# A centromeric RNA-associated protein complex affects germ line development in *Drosophila melanogaster*

**DOI:** 10.1101/2022.02.05.479222

**Authors:** Saskia L. Höcker, Izlem Su Akan, Alexander M. Simon, Kerem Yildirim, Lili A. Kenéz, Ingrid Lohmann, Sylvia Erhardt

**Affiliations:** Karlsruhe Institute of Technology (KIT), Zoological Institute, Fritz-Haber-Weg 4, 76131 Karlsruhe, Germany; Heidelberg University, Centre for Organismal Studies (COS) Heidelberg, 69120 Heidelberg, Germany; Heidelberg University, Centre for Molecular Biology Heidelberg (ZMBH), 69120 Heidelberg, Germany

**Keywords:** centromeres, pericentromeric heterochromatin, satellite repeats, non-coding RNA, RNA-binding proteins, germline, oocyte, development, *Drosophila melanogaster*

## Abstract

In many metazoans, centromeres are embedded in large blocks of highly repetitive (peri-) centromeric heterochromatin from which non-coding RNAs emanate that have been assigned diverse functions in different species. However, little is known about their functional details or regulation. The pericentromere of the X chromosome in *Drosophila melanogaster* contains a multi mega-base array of the 359 bp satellite repeats from the 1.688 family, which is transcribed into a lncRNA (*SAT III* RNA). We performed a *SAT III* RNA pulldown assay and identified a *SAT III* RNA-associated complex of four previously uncharacterized proteins and show that they affect germline development. These factors not only interact with each other and with *SAT III* RNA but also co-regulate each other. RNAi depletion of any of the factors leads to severe defects in the developing germline and sterility. Moreover, we show that the complex plays a crucial role in *SAT III* RNA repression, as RNAi depletion of the factors leads to a drastic increase of SAT III RNA levels. Importantly, genetic reduction of *SAT III* RNA level in the RNAi-depleted flies partially rescued the germ line defects and infertility phenotype. Based on our results we hypothesize that the identified complex functions in the germline to regulate *SAT III* RNA levels, possibly to offset effects of chromatin remodelling taking place in the developing germline.

## Introduction

Centromeric chromatin forms the base for the microtubule fibers of the spindle apparatus to attach and apply force to pull sister chromatids to the opposite poles during anaphase. Although centromeres are usually found at the same chromosome location, their formation is independent of the underlying DNA sequence and, therefore, defined epigenetically. Its determinant is the histone H3-variant CENP-A that is found at a high density at centromeric chromatin, where it replaces canonical H3 in a subset of nucleosomes. In metazoans, centromeric and pericentromeric DNA is usually highly repetitive over megabases of DNA, but low in sequence conservation. To what extent repetitive DNA sequence and unusual chromatin structures are involved in the epigenetic determination of centromeres is still unclear (Mellone & Fachinetti, 2021).

Centromeric and pericentromeric regions are transcribed in all species studied to date and several functions have been proposed for the actual act of transcription at those sides or for the resulting non-coding transcripts (Corless, Hocker et al., 2020). In fission yeast, pericentric transcripts are processed into small interfering RNAs (siRNAs) by the RNA interference pathway, which is important for heterochromatin formation (Volpe, Kidner et al., 2002). In human, mouse, frog and Drosophila, long transcripts can be detected, which have profound functions on cell division, most likely by affecting the integrity of centromeric chromatin, kinetochore formation and cohesion (Blower, 2016, Chen, Zhang et al., 2021, Ferri, Bouzinba-Segard et al., 2009, Ideue, Cho et al., 2014, McNulty, Sullivan et al., 2017, Rosic, Kohler et al., 2014, Wong, Brettingham-Moore et al., 2007). Both depletion and upregulation of (peri)centromeric transcripts can impair mitosis, and upregulation has also been associated with cellular stress, cancer and aging (De Cecco, Criscione et al., 2013, Hedouin, Grillo et al., 2017, Jolly, Metz et al., 2004, Rosic et al., 2014, Swanson, Manning et al., 2013, Ting, Lipson et al., 2011, Valgardsdottir, Chiodi et al., 2005, Valgardsdottir, Chiodi et al., 2008). In humans, (peri)centromeric transcription seems to be partially controlled by the localization of centromeres in the proximity of nucleoli, where centromeric alpha-satellite expression is repressed (Bury, Moodie et al., 2020, Wong et al., 2007). Apart from that, the regulation of (peri)centromeric transcription is not well understood.

In Drosophila, expression of pericentromeric satellite repeats has been well documented, for instance in preblastoderm embryos (Bury et al., 2020, Wong et al., 2007) and germ cells (Wei, Eickbush et al., 2021). In the female gonad, germline stem cells first divide asymmetrically to allow one cell to start differentiating while the other cell retains its stem cell properties. This asymmetry is partially mediated by an asymmetrical distribution of centromere and kinetochore proteins (Dattoli, Carty et al., 2020, Ranjan, Snedeker et al., 2019). The ensuing cystoblast daughter cell undergoes symmetrical divisions, however, without complete abscission, resulting in a 16-cell cyst which will become an egg chamber. One of these cells will develop into the oocyte with heavily compacted chromatin, while the other 15 become nurse cells and start endoreduplicating their genome, including the pericentromeric satellite DNA for the first four endocycles. Subsequently, in endocycle 5, the chromosomes are highly compacted in five DAPI-rich regions representing all major chromosome arms, which has been termed the ‘five-blob stage’. At the onset of endocycle 6, the polytene chromosomes disperse into 32 sister chromosome pairs and continue to endoreduplicate the chromosome arms without further amplifying the (peri)centromeric DNA (Dej & Spradling, 1999). Interestingly, RNA FISH experiments have detected satellite transcripts in endoreduplicating nurse cells (Wei et al., 2021), suggesting that satellite RNAs play a role outside of mitosis. It is, however, unknown whether satellite transcripts contribute to processes such as asymmetric cell division or endoreduplication. It is of note that (peri)centromeres are heavily populated by different classes of transposable elements (TEs), which are also transcribed (Chang, Chavan et al., 2019). To avoid possible damage to the genome, the germ line has developed specialized pathways to transcriptionally silence TEs: the so-called piRNA pathway, which processes piwi-interacting RNAs that target TEs for silencing (Khurana & Theurkauf, 2010). The expression of piRNA precursors is maintained by the interaction of the PolII machinery with heterochromatin components such as the HP1-variant Rhino, via the TFIIA paralog Moonshiner (Andersen, Tirian et al., 2017). However, it has not been established whether other repetitive regions are regulated in a similar manner.

In this study, we focused on the regulation and function of repetitive *SAT III* RNA from the X chromosome of *Drosophila melanogaster* by analysing *SAT III* RNA-interacting proteins. RNA pulldown experiments with *SAT III* transcripts identified a novel nucleolar protein complex that we show to associate with *SAT III* RNA. This complex is highly expressed in the germ line of flies and required for accurate gonadal development, nurse cell chromatin dispersal and fertility. Importantly, we show that the complex is essential for repression of *SAT III* RNA in the germ line and that elevated levels of *SAT III* RNA are at least in part responsible for the observed defects. The complex is involved in controlling *SAT III* RNA levels which in turn allows accurate germ line development in the female reproductive tract.

## Results

### 1. Identification of a SAT III RNA-containing protein complex

The acrocentric X chromosome in *Drosophila melanogaster* contains a large pericentromeric region from which the long non-coding *SAT III* RNA is transcribed in sense and antisense direction. *SAT III* RNA remains nuclear and associates with centromeric and pericentromeric chromatin (Rosic et al., 2014). To identify *SAT III* RNA-interacting proteins, we performed RNA pulldowns from *Drosophila melanogaster* S2 cell lysate, using *in vitro* transcribed *SAT III* RNA that contained 4 copies of the S1m loop, followed by mass spectrometry (Leppek & Stoecklin, 2014) (Fig. **1A** and **S1A**). The experiment was conducted with sense and antisense *SAT III* RNA in separate experiments and in duplicates. *S1m* RNA loops only were used as a control to exclude factors that bind to the loops instead of *SAT III* RNA (Fig. **1B** and **S1B** and **suppl. Tables 1 + 2**). *The SAT III sense* RNA pulldowns were slightly more consistent when compared to the *SAT III antisense* RNA pulldown and we, therefore, initially focused on proteins enriched in the sense pulldown. In total, 72 proteins were enriched in both *SAT III* sense RNA pulldowns, with 36 (50%) ribosomal proteins, 15 (21%) RNA-processing proteins and 6 (8%) uncharacterised proteins of which CG13096 was also highly enriched in the antisense RNA pulldown (Fig. **S1C**). We included three additional proteins (CG1234, CG8545, and CG32344) in our analysis that were only enriched in the first pulldown because a string analysis predicted that they interact with the proteins CG13096 and CG12128 that were enriched in both SAT III sense pulldowns (Fig. **1C** and **S1D**).

**1.**
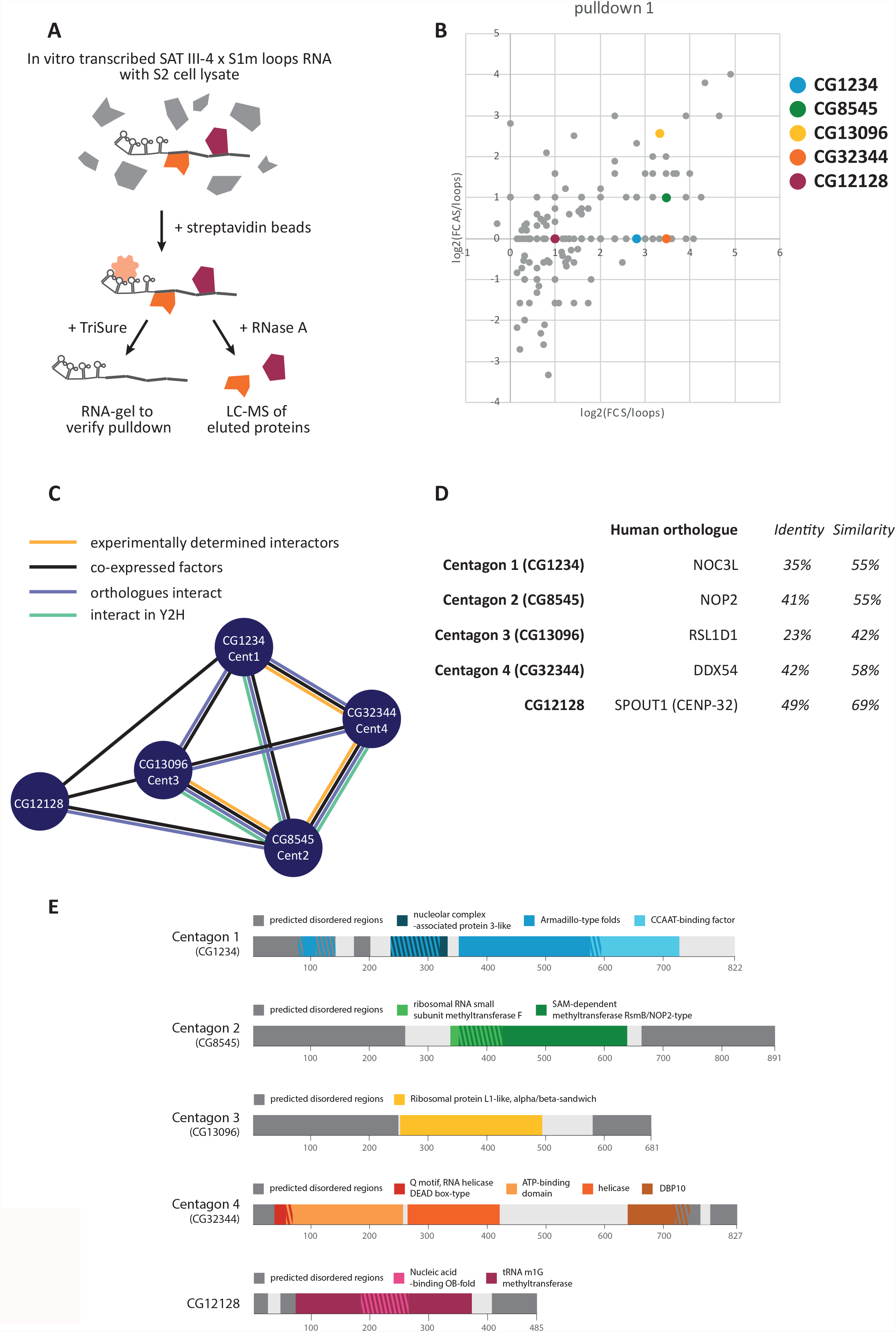
Identification of a SAT III RNA-containing protein complex. **A**. Schematic workflow of the *SAT III* RNA pulldown experiments. *In vitro* transcribed *Sat III* RNAs with 4xS1m loops were incubated with S2 cell lysate and streptavidin beads to pulldown RNA-protein complexes. Approximately 15% of the beads was treated with Trizol for RNA extraction and the rest was incubated with RNase A to elute the *SAT III* RNA-bound proteins, which were subjected to LC-MS. **B**. Enrichment of proteins in the *SAT III* sense and antisense pulldown compared to the only *4xS1m loop* RNA pulldown. Each dot represents a protein and its fold change (FC) in the *SAT III sense* RNA pulldown (x-axis) and *SAT III antisense* RNA pulldown (y-axis) compared to the control on a logarithmic scale. Coloured dots represent members of the Centagon complex and CG12128. A full list of identified proteins can be found in **supplemental table 1. C**. Five uncharacterized proteins found in the *SAT III* RNA pulldown are predicted to form a complex. Data from STRING analysis and our Yeast-Two-Hybrid experiments (see Fig. 2C). **D**. Human orthologues of the five selected factors. Percentages of identical or similar amino acids of the human orthologues to the fly proteins are indicated. **F**. Overview of predicted domains of the candidate proteins according to https://www.ebi.ac.uk/interpro/. Sizes in amino acids are indicated below the protein domains.

The five selected candidates stood out not only for being largely uncharacterized in flies but also for their putative functions based on their human orthologs (Fig. **1D** and **E**) with established links to mitosis or centromeric RNAs. The human orthologue of the putative ribosomal-like protein CG13096 (RSL1D1) was identified in a human *alpha satellite* RNA pulldown (Zhu, Hoong et al., 2018) and the human orthologue of the putative RNA methyltransferase CG12128 (SPOUT1 or CENP-32) was enriched at mitotic spindles and kinetochores, being important for tethering the mitotic spindle to centrosomes and kinetochores (Ohta, Bukowski-Wills et al., 2010, Ohta, Wood et al., 2015). The nucleolar protein CG1234 had been identified to cause shorter mitotic spindles upon knockdown (KD) (Somma, Ceprani et al., 2008), and the human orthologue (NOP2) of the putative RNA-methyltransferase CG8545 was enriched in a human *alpha satellite* RNA pulldown (Zhu et al., 2018). In addition, CG8545 was co-immunoprecipitated with CENP-A in Drosophila cells (unpublished data from our lab). Similarly, the putative DEAD-box RNA helicase CG32344 was found in a CENP-A pulldown of associated factors in Drosophila S2 cells (Barth, Schade et al., 2015). Generally, DEAD-box helicases have been shown to bind *satellite I* ncRNAs in humans contributing to accurate chromosome segregation (Nishimura, Cho et al., 2019). Even though CG12128 is an interesting candidate found in both *SAT III* RNA pulldowns, subsequent experiments showed that it does not share the same phenotypes as the other four candidate proteins and was, therefore, either excluded from our studies or used as a control. The remaining four uncharacterized proteins turned out to be expressed in the gonads and to be important for germline development during subsequent experiments (Fig. **S2** and data below). We named this complex the Centromeric Transcript-Associated Gonadal (**Centagon**) complex, with CG1234 as Centagon 1 (Cent1), CG8545 as Centagon 2 (Cent2), CG13096 as Centagon 3 (Cent3), and CG32344 as Centagon 4 (Cent4) (Fig. **1C-D**).

### 2. *Sat III* RNA interacts with a novel protein complex that exhibits cell cycle-specific localisation patterns

We tested the mutual interactions of the four Centagon proteins with a Yeast-Two-Hybrid assay in triplicates and identified a consistent interaction of Cent2 with Cent, Cent3, and Cent4 (Fig. **2A**, red circles). Empty vectors served as negative control and an interaction of Rdx and Cal1 as positive control (Bade, Pauleau et al., 2014). Furthermore, we validated the binding of *SAT III* RNA to Cent3 using an Electromobility-shift assay (EMSA) (Fig. **2B**). We were able to efficiently purify Cent3-GST in sufficient amounts to perform EMSAs but were unable to do so for the other Centagon proteins. Purified Cent3-GST protein was incubated with *SAT III* and control RNAs at different molar ratios. Addition of Cent3-GST resulted in a band shift of both *SAT III sense* and *antisense* RNA, as well as of the control lncRNA *hsr omega* and an *alpha tubulin* RNA fragment. We concluded from this broad RNA binding spectrum that Cent3 is most likely a bona fide RNA-binding protein with little RNA specificity, at least *in vitro* with purified components. No shifts were observed in control EMSAs with BSA and *SAT III* RNA (Fig. **S1E**). The interaction with the Cent3 subunit demonstrates that the Centagon complex has RNA binding capacity and can directly interact with *SAT III* RNAs.

**2.**
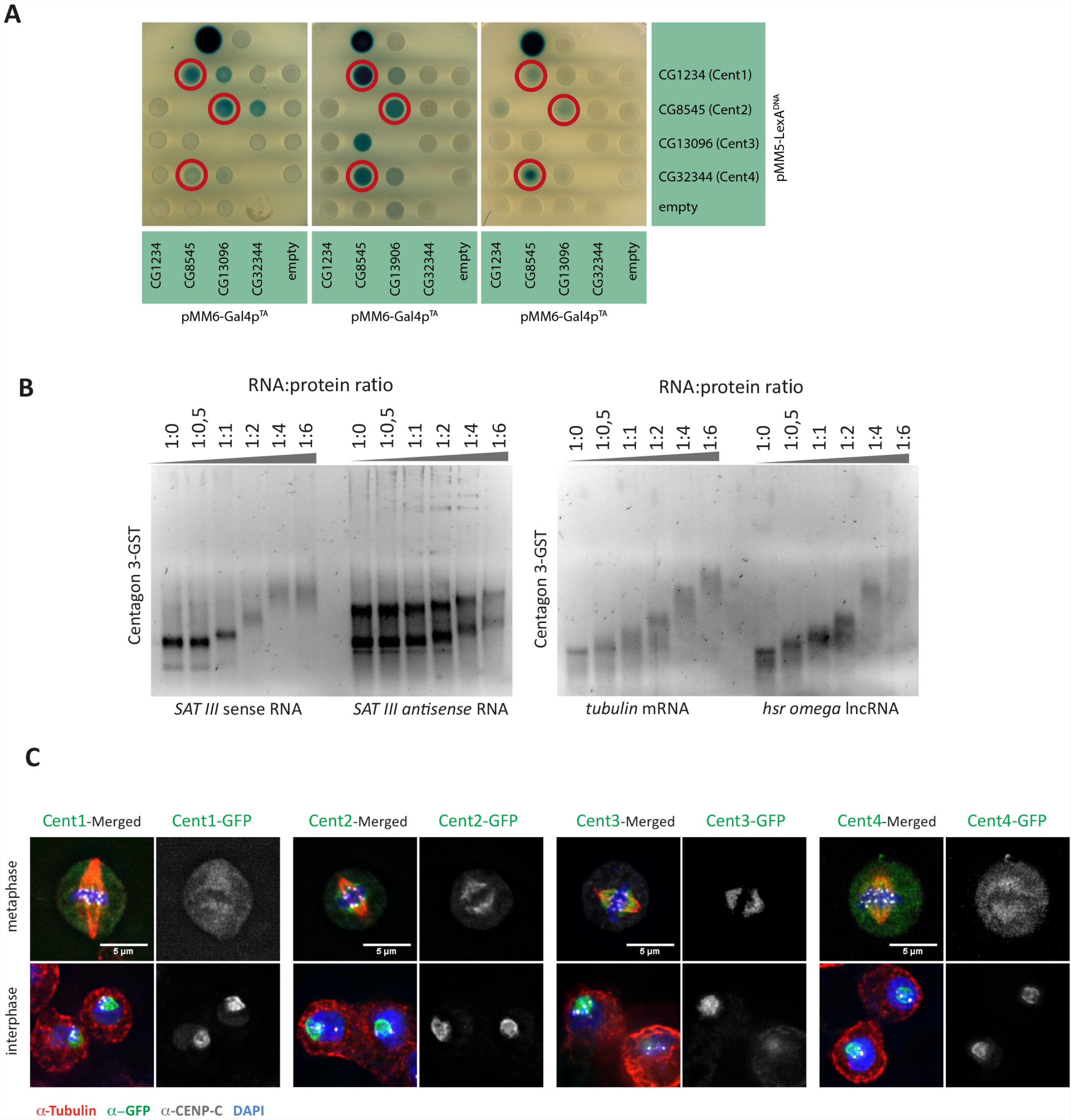
*Sat III* RNA interacts with a novel protein complex that exhibits cell cycle-specific localisation patterns. **A**. Yeast-Two-Hybrid (Y2H) assay with the candidate genes. Three replicates are shown. Only interactions positive in all three assays are encircled in red. The top row depicts the positive (Rdx and Cal1, left) (Bade et al. 2014) and negative (empty plasmids, right) controls. **B**. *SAT III* RNA interacts with Centagon 3 in an electromobility shift assay (EMSA). *In vitro* transcribed RNA was incubated with purified Centagon 3-GST protein in the indicated molar ratios. As RNA controls, the lncRNA *hsr omega* and an *alpha tubulin* RNA fragment were used. **C**. Exogenously GFP-tagged Centagon proteins imaged in metaphase (top row) and interphase (bottom row) S2 cells with the different Centagon proteins in green, anti-tubulin in red, anti-CENP-C in white and DNA in blue. The Centagon complex localises to the nucleolus during interphase and within the spindle area during mitosis. Scale bar 5 μm.

To assess the cellular localization of the Centagon complex, we initially expressed GFP-tagged Centagon proteins in S2 cells and tracked them during the cell cycle (Fig. **2C** and **S3A**). All four proteins show a similar dynamic localization pattern: they are detected at the nucleolus in interphase, overlap with α-tubulin at the microtubule spindle during mitosis and reform nuclear foci in early G1. However, RNAi of the different Centagon complex components did not result in any obvious defects in mitosis, cell cycle regulation or proliferation (data not shown). We, therefore, obtained expression data from the modENCODE Tissue Expression Database (Brown, Boley et al., 2014) and found that all Centagon proteins are most highly expressed in Drosophila ovaries (Fig. **S2**). This suggests that the function of the Centagon complex is tissue-specific and can likely be found in the female germ line.

### 3. The Centagon complex is highly expressed in the germline

In order to assess the localization of Centagon proteins in fly gonads, we produced fly lines with endogenously GFP-tagged Centagon proteins using CRISPR/Cas9 genome editing and imaged larvae and adult fly tissue (Fig. **3A-D**). As expected from their expression profiles (Fig. **S2**), Centagon proteins were detected in the nuclei of both the germline (nurse cells and oocyte) and somatic follicle cells (Fig. **3A**). Co-immunostaining with the nucleolar marker modulo showed that also in ovaries the Centagon proteins localize to the nucleolus in interphase (Fig. **S3B**). We next examined the expression during development and found strong expression already in 3^rd^ instar larval gonads (Fig. **3B** **+ C**). It should be noted here that Centagon 1-GFP and Centagon 4-GFP flies were heterozygous since homozygous stocks were not viable. The weaker GFP signals in those lines may be attributed to their heterozygosity rather than a lower expression compared to Cent2 and Cent3. Importantly, Centagon protein expression is not limited to the gonads, as GFP signal was also observed in larval brain tissue, larval imaginal discs and fat tissue (Fig. **3B** **+ C**). In the germarium, Centagon proteins are highly expressed in the germ stem cells (GSCs, marked by pMad immunostaining) and first cystoblasts, but undetectable in cells derived from subsequent divisions (Fig. **3D**). The Centagon complex is also expressed in the accompanying somatic cells. Only at egg chamber stage 1, expression increases again in the germ cells, indicating a differential Centagon expression pattern during early germ cell divisions.

To confirm that the Centagon complex also binds *SAT III* RNA *in vivo* and *in ovary*, we performed RNA-Immunoprecipitation (RIP) with ovary lysate from the two homozygous lines of the endogenously tagged Centagons, Cent2 and Cent3, and analysed the *SAT III* RNA levels by qPCR. We detected a significant enrichment of *SAT III* RNA in the Cent2-IP compared to the control *GAPDH* mRNA (Fig. **3E** **+ S3E**). *SAT III* RNA was also enriched in the Cent3 RIP but only two out of three repeats showed a strong enrichment (Fig. **3F** **+ S3E**). This result suggests that also in ovaries, the Centagon complex binds *SAT III* RNA.

**3.**
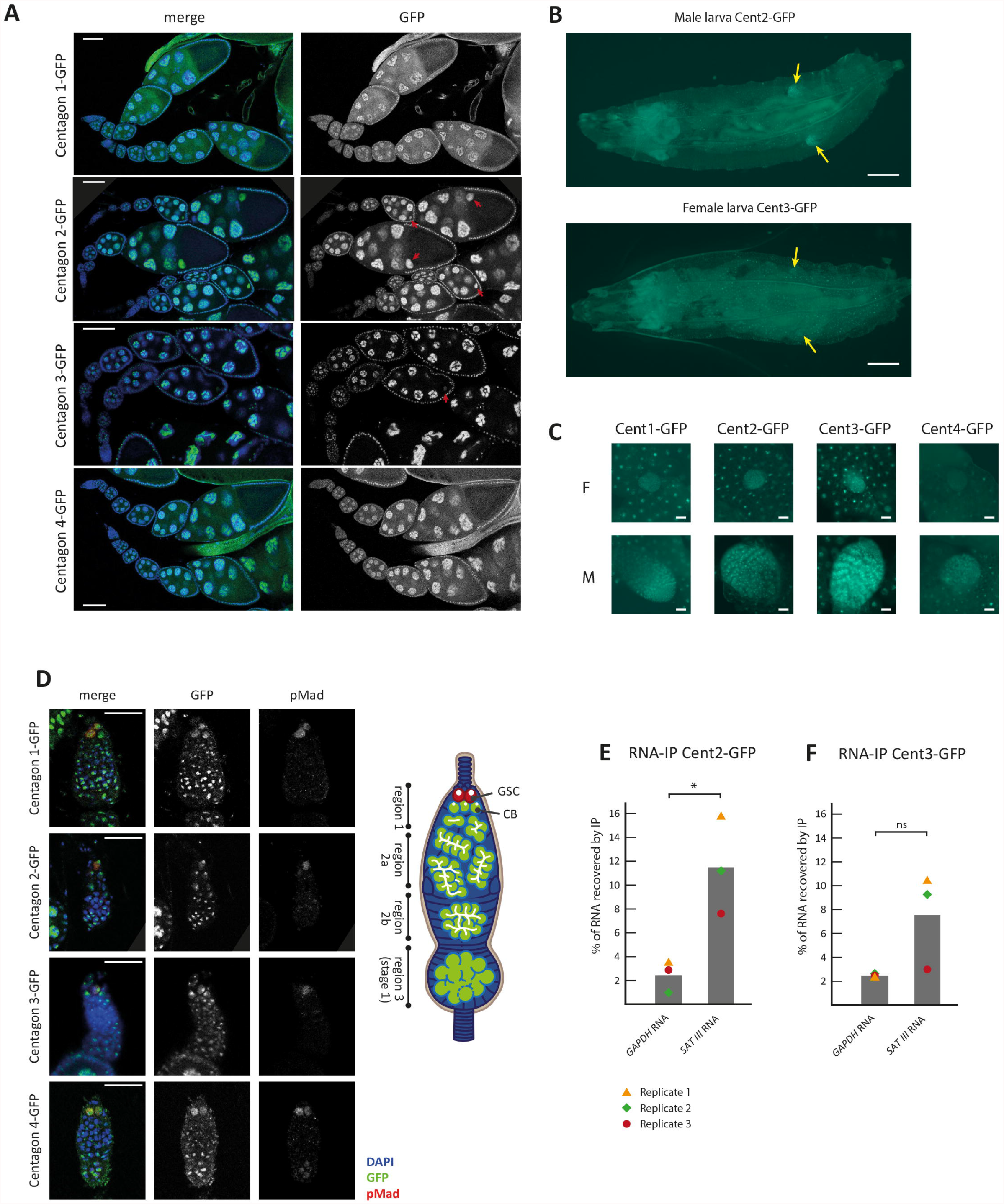
The Centagon complex is highly expressed in the germline. **A**. Ovarioles of flies with endogenous GFP-tagged Centagon proteins. The Centagon complex localises to the nuclei of oocytes (red arrows), nurse cells and follicle cells of the Drosophila ovary. Scale bar 50 μm. **B**. Live imaging of larvae with endogenous GFP-tagged Centagon proteins, the location of the gonads is indicated by arrows. Scale bar 0,5 mm. **C**. Female (F) and male (M) dissected gonads from 3^rd^ instar larvae surrounded by fat tissue with endogenous GFP-tagged Centagon proteins. Scale bar 50 μm. **D**. Germaria of flies with endogenous GFP-tagged Centagon proteins (green), co-immunostained with the germ stem cell marker pMad (red) and DAPI (blue). For easier orientation, a model of the germarium has been included on the right and some of the immunofluorescent images were rotated on top of a grey background. Scale bar 20 μm. **E+F**. RNA-IP with ovary lysate of the homozygously tagged Cent2-GFP (**E**) or Cent3-GFP (**F**) flies, followed by qPCR to measure the levels of associated *SAT III* RNA and *GAPDH* mRNA as control. RNA levels of input and IP samples were compared to calculate the percentage of recovered RNA in the IP.

### 4. The Centagon complex is important for oogenesis

To study the germ line function of the Centagon complex we next performed knockdown experiments of the Centagon members in ovaries. To do so, we crossed TRiP RNAi fly lines targeting Centagon transcripts with two ovary-specific Gal4 driver lines: Maternal Triple driver (MTD-Gal4) and Mat67.15-Gal4 (Grieder, de Cuevas et al., 2000, Mazzalupo & Cooley, 2006, Staller, Yan et al., 2013). MTD-Gal4 is expressed already in GSCs and Mat67.15-Gal4 starts expression in early egg chambers (Fig. **S3C**). We dissected the ovaries of the resulting female offspring and observed an obvious size difference between control and all Centagon KD ovaries. In contrast, KD of CG12128, which we used as control, did not affect ovarian development (Fig. **S3D**). KD of the Centagon members during early germ cell development (with MTD-Gal4) led to more severe phenotypes with strongly underdeveloped ovaries compared to KD at later stages (with Mat67.15-Gal4) suggesting a function during early germ line development. The efficiency of the Mat67.15-induced KD was monitored by qPCR (Fig. **4A**). Not only were the mRNA levels of the Centagon members significantly reduced when depleted by RNAi, upon KD of one member, levels of the other three Centagon members were significantly upregulated, emphasizing the existence of a functional complex and suggesting co-regulation and interdependence of the Centagon complex components. No qPCR data for the MTD-induced KD efficiency is available because there was too little tissue left after knockdown. However, since the phenotype was consistent with the one from the Mat67.15 induced KD, we are confident that the KD was also highly efficient. Immunofluorescent imaging showed that in the early MTD-induced Centagon KD ovaries, almost no vasa-positive germ cells were left in the germaria and only very few immature egg chambers were observed (Fig. **4B**, white arrows). Interestingly, this was not due to a loss of GSCs, since many pMad-positive cells were left in the stem-cell niche. However, we observed aberrant pMad expression at the germ cell cyst stage (red arrows), which was significantly higher in the Centagon 1 KD ovaries compared to the control (Fig. **4C**). This result indicates a possible differentiation defect of germ cells upon Centagon depletion with cells maintaining a stem cell-like potential longer than under control conditions.

**4.**
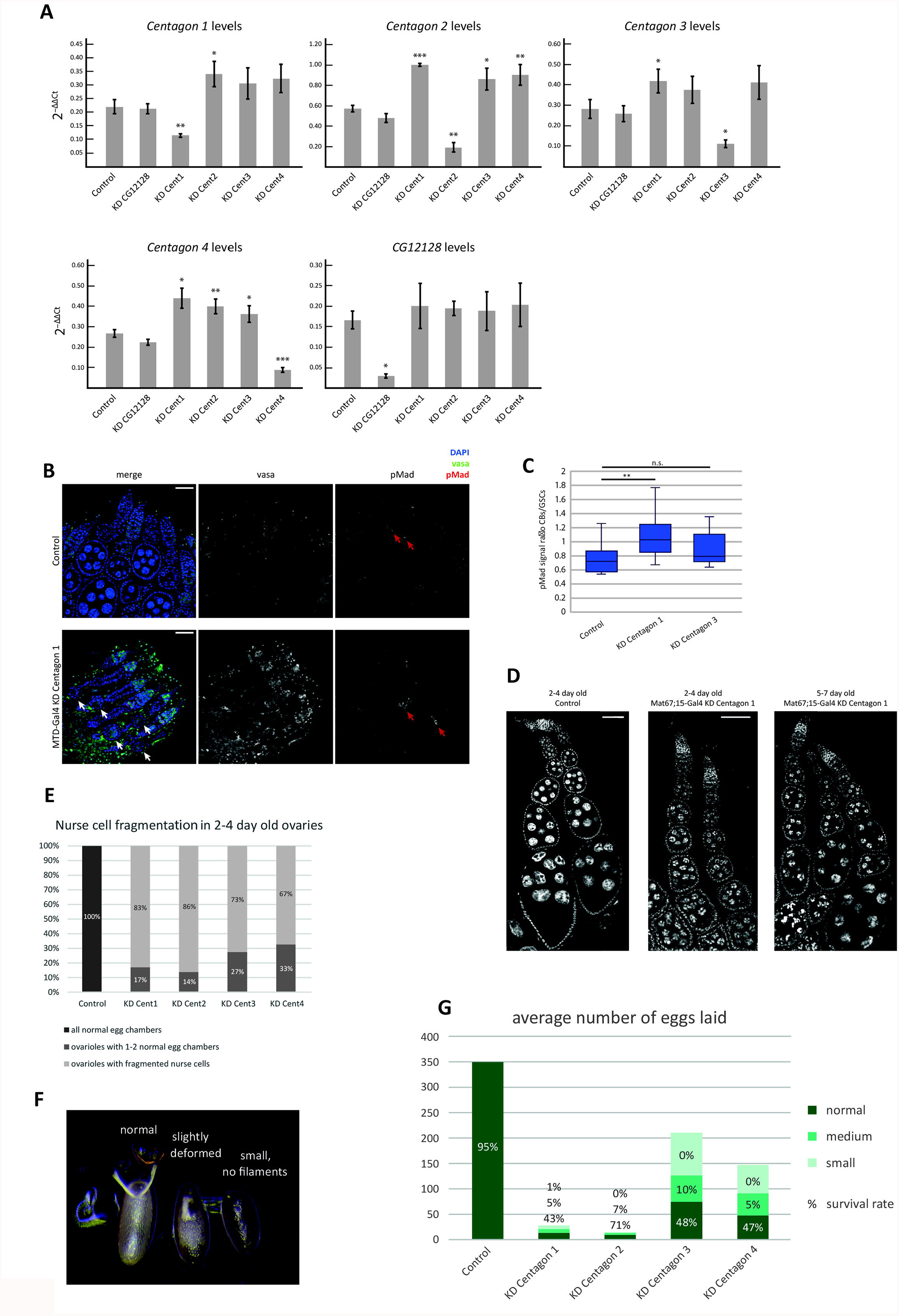
The Centagon complex is important for oogenesis. **A**. qPCR of Centagon RNA levels in ovaries after KD with the Mat67.15-Gal4 driver. RNA levels are normalized to housekeeping genes. N=3. Student t-test, * p ≤ 0.05, ** p ≤ 0.01, *** p ≤ 0.001. **B**. 0-1-day old ovaries of control and MTD-Gal4-induced Cen1 KD flies. Co-immunostaining with the germ cell marker vasa (green) and the germ stem cell marker pMad (red), and DAPI (blue). White arrows point towards emerging egg chambers in the KD ovaries. Red arrows point towards aberrant pMad signal. Scale bar 25 μm. **C**. Quantification of the difference in pMad signal per cell in cystoblasts vs. GSCs as shown in B. Student t-test, ** p ≤ 0.01. **D**. DAPI staining of control and Mat67.15-Gal4-induced Centagon KD ovarioles derived from flies of different ages. For easier orientation, some images were rotated on top of a grey background. Scale bar 50 μm. **E**. Quantification of the phenotype shown in D. Egg chambers with and without the five-blob chromatin phenotype (after egg chamber stage 5) were counted within an ovariole. **F**. Eggs (F2) derived from Mat67.15-Gal4 driver-induced Centagon KD flies (F1). The observed phenotypes were classified into the categories normal, medium (slightly deformed) and small (also without dorsal filaments). **G**. Number of eggs laid overnight by Mat67.15-Gal4-induced Centagon KD flies. The egg phenotypes are indicated by colour and the survival rate of each egg category is indicated by percentage.

When the Centagon members were knocked down in early egg chambers (with Mat67.15-Gal4), egg chambers ceased to develop and displayed abnormally fragmented nurse cell nuclei that were reminiscent of the 5-blob phenotype (Fig. **4D** **+ E**) (Dej & Spradling, 1999). In these ovarioles, most egg chambers degenerated before reaching a mature egg stage. The phenotype was strongest in young females (2-4 days). In older females (5-7 days), nurse cells were less fragmented and more egg chambers developed (right panel), indicating some adjustment of the phenotype with age. However, the eggs laid by both young and old females were smaller with no dorsal filaments, which we categorized into three groups: normal, medium (slightly deformed) and small (without dorsal filaments) (Fig. **4F**). A survival assay showed that KD of the Centagon members in the ovary leads up to a 20-fold reduction of eggs laid overnight (KD of Cent2) with most of them being deformed. Unsurprisingly, there is a clear correlation between egg phenotype severity and the hatching rate (survival rate) of these eggs after 24 hours (Fig. **4G**). We concluded that the Centagon complex is both important during early germ cell divisions and for egg chamber development.

As the Centagon complex is also expressed in somatic ovary cells and germ cells of the testis, we also conducted KD experiments in these cell types. Depletion in the ovary follicle cells resulted in compound egg chambers, follicle cell gaps and, in case of depletion in follicle stem cells, to loss of the entire ovary (Fig. **S4A** **+ B**). It is important to note that even though the expression profiles for Centagon components in adult testis are not as high in the FlyAtlas Anatomical Expression (Chintapalli, Wang et al., 2007) and modENCODE Tissue Expression Data sets (Brown et al., 2014) compared to adult ovaries, we detected strong GFP signals in the developing testis of endogenously tagged flies and a strong male germ line phenotype upon depletion. Centagon depletion in the testis led to smaller, deformed testes with a misplaced hub and no sperm formation (Fig. **S4C**). Therefore, we concluded that the Centagon complex is important for the development of somatic and germ cell development in the male and female germ line of *Drosophila melanogaster*.

### 5. *SAT III* RNA levels are repressed by the Centagon complex in ovaries

As the Centagon complex was initially identified as a *SAT III* RNA-associated complex, we next investigated if the depletion of Centagon members has an effect on *SAT III* RNA levels and localization. First, we performed *SAT III* RNA FISH in ovaries to determine the *SAT III* RNA localization in w^1118^ control flies (Fig. **5A**). In w^1118^ ovarioles the *SAT III* RNA localized in one to several foci in nurse cell nuclei as well as inside the oocyte nucleus (see close-up). To test the specificity of the FISH signal, we used the Zhr^1^ fly line which carries a chromosomal translocation that removed the majority of the pericentric SAT III DNA region (Ferri et al., 2009, Sawamura, Yamamoto et al., 1993). We tested both *SAT III sense* and *antisense* RNA FISH, but observed a more specific signal with the antisense probes: Although the signals of *SAT III* sense and antisense probes showed similar localizations in ovarioles, some *SAT III sense* RNA FISH signal was also detectable in Zhr^1^ flies (Fig. **S5A**), indicating that there are a few SAT III repeats remaining in Zhr^1^ flies or that the SAT III sense probe may also detect other satellite repeats. Therefore, further FISH experiments were conducted with SAT III antisense probes. Indeed, Zhr^1^ flies did not show any of the *SAT III* RNA FISH foci that we observed in w^1118^ ovarioles (Fig. **5A**). To ensure that the observed signal in control flies was indeed RNA and not DNA, which are both mostly lacking in Zhr^1^ flies, we treated the ovaries with Rnase A. All *SAT III* RNA foci in w^1118^ ovaries disappear after treatment with RNase A, indicating the specific hybridization of SAT III probes to RNA (Fig. **S5B**). Importantly, the *SAT III* RNA FISH signal increased drastically in Mat67.15-induced Centagon KD ovaries (Fig. **5A** **+ B**). Although the Centagon members were mostly identified in the *SAT III sense* RNA pulldown, the RNA FISH experiments show that *SAT III sense* and *antisense* transcripts are both upregulated after Centagon KD. This increase was further confirmed by qPCR, where we measured an up to 50-fold increase of *SAT III* RNA levels in Centagon KD ovaries compared to the control (Fig. **5C**). Furthermore, the spatial resolution of the *SAT III* RNA FISH revealed that *SAT III* RNA levels were specifically increased in egg chambers with fragmented nurse cells. To quantify this effect, we measured *SAT III* RNA levels in KD ovaries of 5-7-day old females with both fragmented and normal nurse cell chromatin and found a highly significant increase of *SAT III* RNA signals in nurse cells with fragmented chromatin (Fig. **5D** and **E**). Comparable results were obtained with the MTD-induced KD where *SAT III* RNA FISH signal increased in KD ovaries (Fig. **S5C**). This indicates that the Centagon complex controls *SAT III* RNA levels and that an increase of *SAT III* RNA levels correlates with fragmented nurse cell nuclei in Centagon KD ovaries.

**5.**
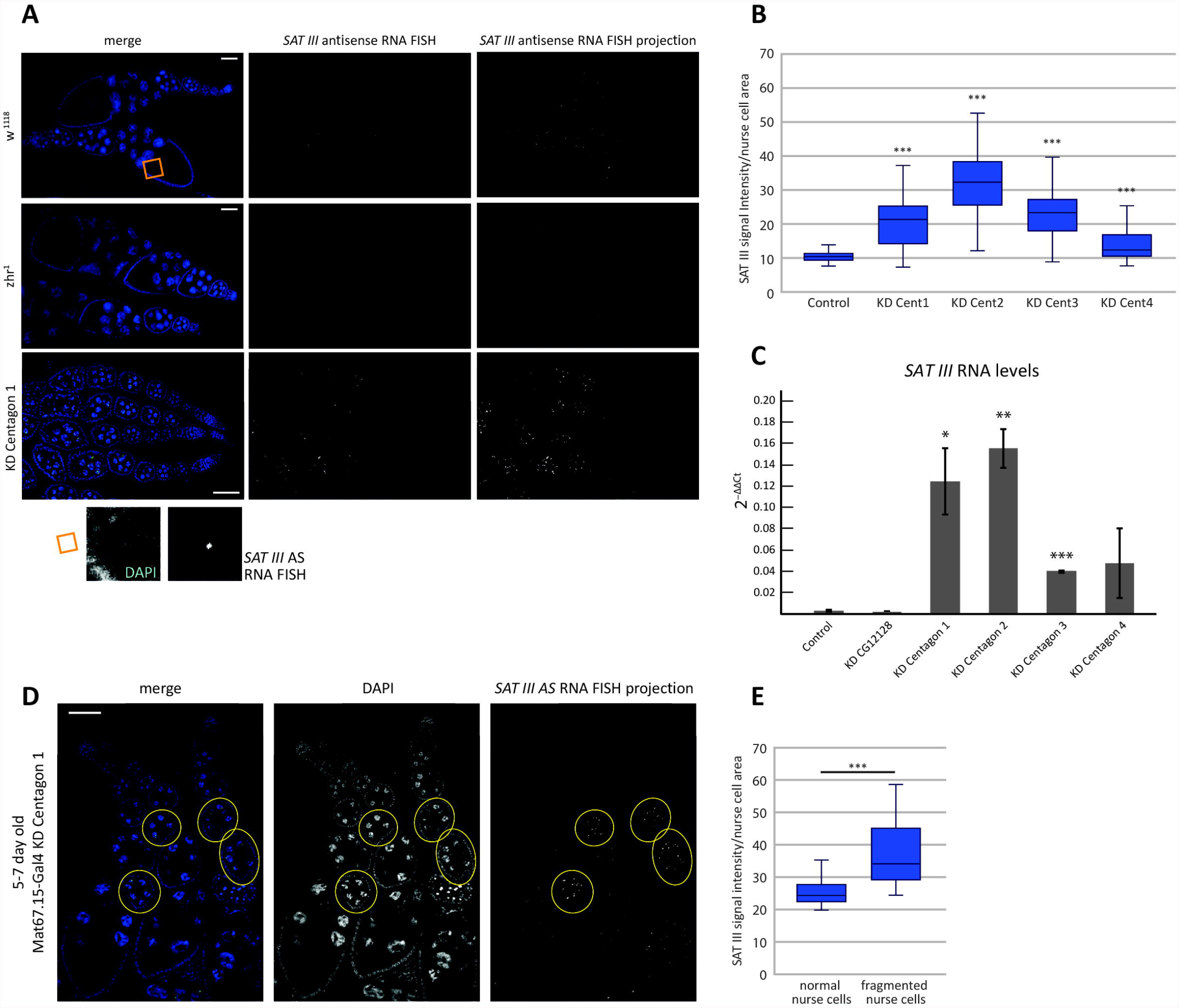
*SAT III* RNA levels are repressed by the Centagon complex in ovaries. **A**. *SAT III antisense* RNA FISH (green) in w^1118^, zhr^1^ and Mat67.15-Gal4-induced Cen1 KD ovaries, stained with DAPI (blue). *SAT III* RNA signals are depicted both in one z-slice (middle) and as z-stack projection (right). The orange square is enlarged on top and shows the oocyte nucleus. For easier orientation, some images were rotated on top of a grey background. Scale bar 50 μm. **B**. Quantification of the *SAT III* RNA FISH signal in A. The *SAT III* RNA FISH signal was measured per egg chamber and normalized to the egg chamber size (measured area). Student t-test *** p ≤ 0.001. **C**. qPCR of ovary *SAT III* RNA levels in Mat67.15-Gal4-induced Centagon KD flies. RNA levels are normalized to housekeeping genes. N=3Student t-test, * p ≤ 0.05, ** p ≤ 0.01, *** p ≤ 0.001. **D**. *SAT III* RNA FISH (green) in 5-7-day old Mat67.15-Gal4-induced Cen1 KD ovaries, stained with DAPI (blue). The *SAT III* RNA signal is depicted as z-stack projection. Egg chambers with aberrant nurse cell fragmentation (5-blob phenotype) are encircled. For easier orientation, some images were rotated on top of a grey background. Scale bar 50 μm. **E**. Quantification of the *SAT III* RNA FISH signal in C per egg chamber (see B). Statistical difference between egg chambers with normal and fragmented nurse cells were determined using a student t-test, *** p ≤ 0.001.

### 6. *SAT III* RNA reduction partially rescues the Centagon KD phenotypes

To evaluate the causal relationship of high *SAT III* RNA levels and nurse cell fragmentation, we genetically lowered the *SAT III* RNA level by crossing the Centagon 1 RNAi-line and Mat67.15-Gal4 driver line with the Zhr^1^ fly line. When Centagon 1 was depleted in ovaries of Zhr^1^, nurse cell fragmentation was significantly reduced (Fig. **6A + B**). Furthermore, females with Centagon 1 depletion and the Zhr^1^ X chromosomes laid as many eggs as the control w^1118^ females (Fig. **6C**). However, many of these eggs were deformed and had a lower survival rate. It should be noted that the KD efficiency of Centagon 1 was slightly lower in Zhr^1^ flies (Fig. **6D**), which could be partially responsible for the reduced phenotype we observed. Additionally, *SAT III* RNA levels were assessed by qPCR (Fig. **6E**), which indeed remained low in Zhr^1^ flies. We concluded that the effect observed after Centagon KD is at least partially caused by high *SAT III* RNA levels resulting from the lack of repressive function of the Centagon complex. In conclusion, we identified a complex that regulates (peri)centromeric *SAT III* transcript levels, which is essential for normal somatic and germ cell development in the female gonad and may have additional functions in the male germline and potentially other tissues where we detected high levels of the Centagon complex.

**6.**
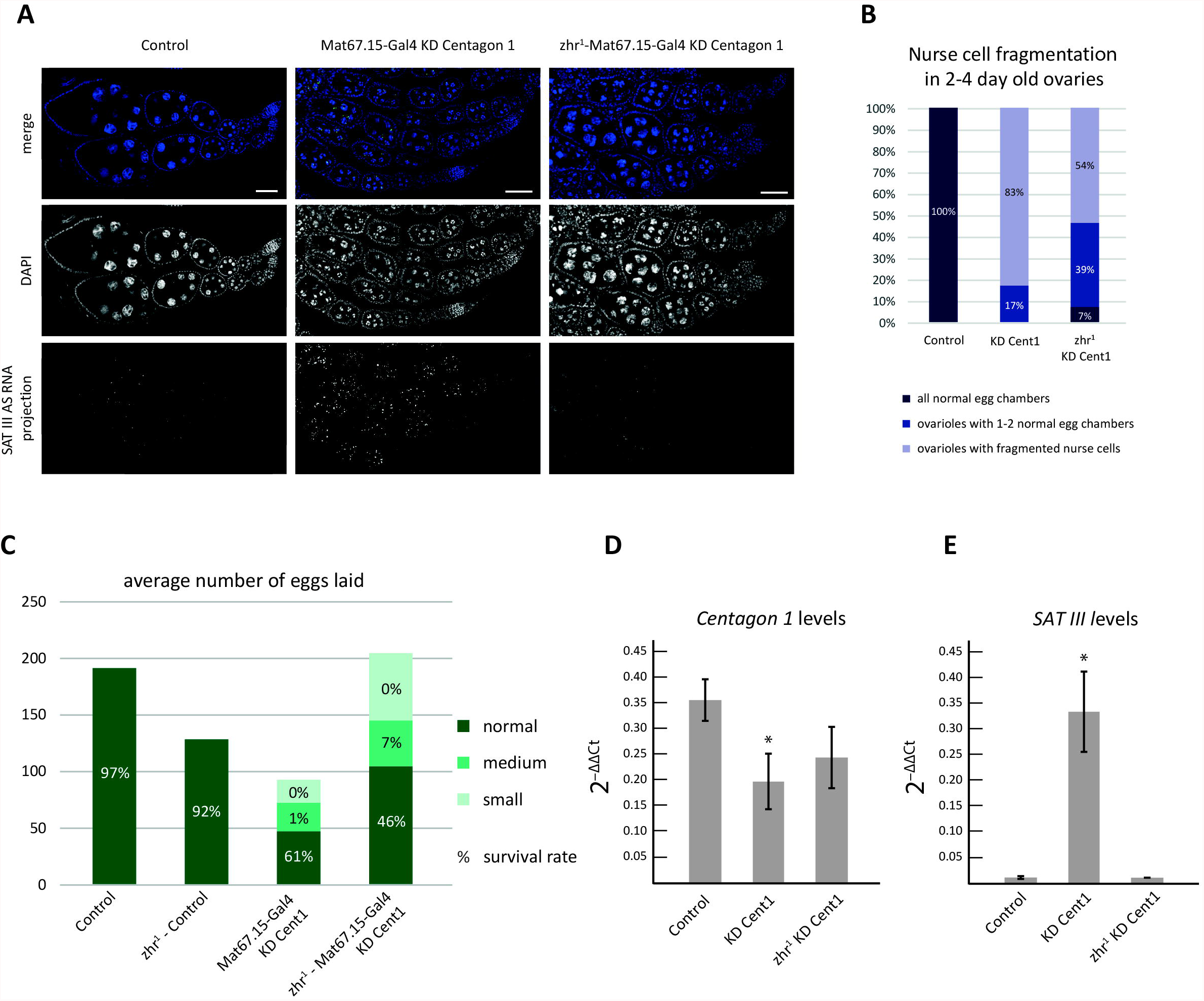
*SAT III* RNA reduction partially rescues the Centagon KD phenotypes. **A**. *SAT III* RNA FISH (green) in control, Cen1 KD and Zhr^1^;Cen1 KD ovaries, stained with DAPI (blue). The *SAT III* RNA signal is depicted as z-stack projection. Rotated pictures are on top of a grey background. Scale bar 50 μm. **B**. Quantification of egg chambers with fragmented nurse cells as in 4E. **C**. Number of eggs laid overnight by Cen1 KD flies with and without SAT III. The egg phenotypes are indicated by colour and the survival rate of each egg category is indicated by a percentage. **D+E**. RNA levels of *Cent1* (D) and *SAT III* (E) in control, Cent1 KD and Zhr^1^;Cent1 KD ovaries measured by qPCR. RNA levels are normalized to housekeeping genes. N=3. Student t-test, * p ≤ 0.05.

## Discussion

There are longstanding questions as to what extent repetitive regions of the genome are functionally relevant and whether transcription or the transcripts themselves are important for genome regulation and cellular functions. In addition, how these transcripts are regulated is not well understood. We focused on the long non-coding RNA *SAT III* from the repetitive (peri-)centromeric region of the X chromosome of *Drosophila melanogaster* and show that *SAT III* RNA is regulated by a novel RNA-binding complex of at least four different proteins that we termed ‘Centromeric Transcript-Associated Gonadal’ (Centagon) complex. We show here that the Centagon complex represses *SAT III* RNA levels in the female gonad and influences germ line development and fertility.

Previously, *SAT III* RNA has been shown to promote correct chromosome segregation during mitosis since depletion of *SAT III* RNA or the genomic locus causes segregation defects in Drosophila cells and embryos (Rosic et al., 2014). Here we show that elevated *SAT III* RNA levels are also detrimental for gonadal cells *in vivo*: The Centagon complex controls *SAT III* RNA levels in the Drosophila gonad and depletion of complex members resulted in an up to 50-fold increase of *SAT III* transcript levels accompanied by massive defects in egg chamber maturation. Importantly, reducing the *SAT III* RNA level in Centagon-depleted ovaries partially rescues the phenotypes. We therefore conclude that accurate levels of *SAT III* RNA are fundamental to cell survival and development, and tightly regulated. Interestingly, the Centagon complex is important in both actively dividing early germ cells and in postmitotic nurse cells during egg chamber maturation (Fig. **4**) indicating an effect of *SAT III* RNA upregulation outside of mitosis. Depletion of the Centagon complex in the germarium during early germ cell divisions led to empty ovarioles and aberrant expression of the germ stem cell marker pMad. This indicates that the Centagon complex is important for correct germ cell differentiation. Interestingly, the Centagon proteins were amongst 646 factors from a transcriptome-wide RNAi screen that aimed to identify networks controlling stem cell self-renewal and differentiation in female gonads (Sanchez, Teixeira et al., 2016).

Because the depletion of Centagon components in S2 cells did not result in obvious defects, its role in regulating *SAT III* levels might be restricted to certain stages and tissues during development as suggested by the high expression levels of endogenously tagged Centagon components in larval tissue. Centagon levels seem particularly high in tissues that undergo changes in their developmental potential such as the germline and imaginal discs. This may include changes in heterochromatin and strict regulation of transcripts from repetitive elements by the nucleolar Centagon complex. This is supported by previous reports that the localization of centromeres to an intact nucleolus is important for heterochromatin formation and transcriptional silencing of repetitive elements in Drosophila and human cells (Bury et al., 2020, Padeken, Mendiburo et al., 2013).

It is important to note that the Centagon complex components have been implicated in ribosome biogenesis: The yeast orthologue of Cent1 is involved in 60S ribosomal subunit maturation (Milkereit, Gadal et al., 2001), while Cent2 as a putative RNA methyltransferase may be involved in rRNA maturation as has been documented for the nucleolar RNA methyltransferase fibrillarin (Tollervey, Lehtonen et al., 1993). KD of Cent3 leads to accumulation of aberrant rRNA intermediates (Sanchez et al., 2016) and DEAD-box RNA helicases like Cent4 are often part of large complexes such as ribosomes or spliceosomes (Linder & Jankowsky, 2011). It is not known whether defects in ribosome biogenesis also affect *SAT III* transcript levels or vice versa, however the SAT III locus on the X chromosome is immediately adjacent to the rDNA locus and a deregulated chromatin state of one of the loci may directly or indirectly affect the other. Furthermore, there seems to be a strong connection between rRNA synthesis and GSC differentiation (Neumuller, Betschinger et al., 2008, Sanchez et al., 2016, Zhang, Shalaby et al., 2014). However, the fact that we did not observe any major defects in S2 cells and no obvious structural defects in the nucleolar organization argues against a major functional importance of the Centagon complex in ribosome biogenesis or nucleolar integrity. More work is needed to fully understand the precise relationship of centromeric transcript and the nucleolus, especially in specialized tissues like the germ line.

KD of the Centagon members in early egg chambers resulted in egg chamber maturation defects, visible by an extended fragmented (5-blob) chromatin state in nurse cells, which remained small in size and eventually degenerated. A clear connection of the observed phenotypes with elevated *SAT III* RNA levels could be established: the simultaneous depletion of Centagon members and *SAT III* RNA reduced the nurse cell chromatin fragmentation phenotype and led to a higher number of eggs. Although the nurse cells do not undergo mitosis, they are subjected to a unique endocycle which requires a mitosis-like step to disperse the nurse cell chromatin after endocycle 5 (Dej & Spradling, 1999), which is apparently partially blocked by high *SAT III* RNA levels. Interestingly, some Centagon orthologues have been implicated in cell cycle functions: Cent1 orthologues in human and yeast and the human Cent2 orthologue promote S phase and replication initiation (Cheung, Amin et al., 2019, Johmura, Osada et al., 2008, Wang, Wang et al., 2020, Zhang, Yu et al., 2002) and the human orthologue of Cent3 promotes proliferation (Ma, Chang et al., 2008).

Germ cells are highly specialized cells in an organism that need to prepare the genome for transitioning from a fully differentiated germ cell to being part of the totipotent zygote after fertilization. This is reflected in massive chromatin remodelling events that must take place during early development to remove highly specialized chromatin organization from the sperm and egg to become a zygote. These early embryos are defined by a largely unconstrained genome conformations until the maternal to zygotic transition when the genome becomes ordered and structured into domains (Hug & Vaquerizas, 2018). We propose that an important function of the novel SAT III silencing mechanism we describe is to keep the transcriptional activation of repetitive elements under control during chromatin remodelling processes in the germ line to maintain genome integrity. This has been well described for the Drosophila germline, where transposable elements are transcriptionally silenced through heterochromatin formation mediated by the PIWI-interacting RNA pathway to protect the genome (Batki, Schnabl et al., 2019, Sienski, Donertas et al., 2012). We envision a similar scenario for repetitive regions that need to be highly controlled by a repressive complex in the germ line, in this case, Centagon. The complex is specifically expressed in the germ line and in tissues that undergo changes in their developmental potential, perhaps to contribute to satellite repeat silencing during chromatin remodelling phases that occur during these developmental stages. Why too much or too little *SAT III* RNA causes developmental defects is still unclear but by identifying the Centagon complex, we are one step closer in understanding *SAT III* RNA regulation and its specific role in germ line development.

## Materials and Methods

### Cloning

Full-length Centagon coding regions were cloned into a modified pMT/V5 vector (Invitrogen) or pLAPcopia-GFP (Erhardt, Mellone et al., 2008) for imaging S2 cells or pGEX for bacterial expression of recombinant proteins; SAT III repeats were cloned into pSP73-4xS1m (Leppek & Stoecklin, 2014, Rosic et al., 2014). All plasmid sequences were verified by Sanger sequencing. For the knock-in transgenic flies, endogenous Centagon loci were N-terminally (CG1234) or C-terminally (CG8545, CG32344) GFP tagged. The guide RNAs were designed using CCTOP (https://cctop.cos.uni-heidelberg.de/index.html (Stemmer, Thumberger et al., 2015). The cloning and injection were performed by Qidong Fungene Biotechnology Co.Ltd. (http://www.fungene.tech). For yeast two hybrid studies, full length Centagon 1-4 and CG12128 were cloned into pMM5 and pMM6 plasmids as described for the controls CAL1 and Rdx in (Bade et al., 2014).

### RNA extraction

RNA from S2 cell pellets was purified using equivalent amount of Trizol (Invitrogen) or TriSure (Meridian Bioscience). Dissected and snap frozen ovaries were supplemented with Trizol (or TriSure) to a total of 100 μl. The tissue was homogenized and the samples were centrifuged at 12.000 x g at 4°C for 10 min. RNA was extracted from the supernatant according to the manufacturer’s protocol. RNA pellets were resuspended in RNase-free water according to the pellet size (20-100 µl).

### gDNA digestion of ovary RNA

10 μg ovary RNA was digested with Turbo DNase (Thermo Fisher Scientific) at 37°C for 20 min. For *SAT III* RNA qPCR from ovaries an additional gDNA digestion step was performed to eliminate DNA contamination. The reaction was mixed with 170 μl H_2_O as well as 20 μl NaAC (3M, pH 5.2) on ice, 220 μl Phenol-Chloroform-Isoamyl added, vortexed, and centrifuged at 16.000 x g for 5 min at RT. The upper phase was supplemented with 200 μl chloroform, vortexed and centrifuged at 16.000 x g 5 min at RT. The upper phase was mixed with 200 μl Isopropanol and 1 μl GlycoBlue (Invitrogen) at -80°C for at least 1 h. The sample was centrifuged and the pellet washed with 75% ethanol and again centrifuged, air-dry and the pellet resuspended in 15 μl H_2_O.

### Reverse Transcription and qPCR

To synthesize cDNA from 1 μg RNA the Quantitect kit (Qiagen) was used according to manufacturer’s instructions. qPCRs were performed using LightCycler® 480 SYBR Green I Master Mix (2X) (Roche) with one reaction containing: 7,5 μl 2x Sybrgreen, 1 μl diluted cDNA, 1 μl each of forward and reverse primers (10 μM) and 4,5 μl H_2_O. The following program was used on the LightCycler® 480: 10 min at 95°C, followed by 40 cycles of 15 sec. 95°C and 1 min at 55°C.

### RNA gel electrophoresis

RNA from the RNA-pulldown experiment was run on a denaturing MOPS-formaldehyde 1% agarose gel in 1/5 volume RNA loading buffer (50% (v/v) glycerol, 1 mM Na_2_EDTA, 0.4% (v/v) bromophenol blue, 40 μg/ml ethidium bromide) and 2x the volume of RNA including loading buffer of RNA sample buffer (0,65x MOPS, 65% (v/v) formamide, 8,5% (v/v) formaldehyde). The RNA was heated at 65°C for 10 min to unfold, immediately placed on ice and loading and separated at 70 V in 1x MOPS (20 mM MOPS pH 7.0, 2 mM sodium acetate, 1 mM EDTA).

### RNA Electromobility Shift Assay (EMSA)

*Recombinant protein purification*. GST-tagged protein in BL21 bacteria were grown to an OD 0.6 density and induced with 0.3 mM IPTG at 25°C for 16 h. Cells were harvested, washed with PBS, incubated in lysis buffer (500 mM NaCl, 0.1% (v/v) NP-40, 2mM PMSF, 1 μg/ml Aprotinin, 1 μg/ml Leupeptin-Hemisulfate, 1 mg/ml Pepstatin, 1 mM DTT in 1x PBS) at 4°C and homogenized in two cycles with the Avestin Emulsiflex and centrifuged at 20.000 rpm at 4°C for 30 min. The lysate was filtered and loaded onto the ÄKTA GST column chromatography system, washes with 150 mM and 100 mM NaCl and eluted with 30 mM Glutathione in PBS, pH 9.0. The eluate was loaded onto a PD-10 desalting column (GE Healthcare) and eluted with PBS to remove salt. Further concentration was performed with a 10K centricon tube (Amnion).

Equal amounts of *in vitro* transcribed RNA (T7 MegaScript kit, Invitrogen) were incubated with increasing amounts of purified recombinant protein. RNA was heated to 68°C for 10 min and incubating 10 min at RT before placing on ice. The RNA and protein were cooled down for 30 min on ice and samples loaded on a non-denaturing 1% agarose gel in TAE buffer (40mM Tris, 20mM Acetate, 1mM EDTA) and run at 120 V followed by a 20 min ethidium bromide bath (RNase-free TAE).

### Yeast Two Hybrid

All full length Centagon sequences were cloned into the pMM5-LexA^DNA^ and pMM6-Gal4p^TA^ vectors (Bade et al., 2014). A combination of one pMM5 and one pMM6 plasmid was transformed into competent yeastS SGY37VIII cells (Knop, Siegers et al., 1999). The YTH was performed as described previously (Bade et al., 2014). In short, interactions were judged based on the activity of ® -galactosidase that results in the conversion of X-Gal (5-bromo-4-chloro-3-indolyl-b-D-galactosidase) into a blue dye.

### RNA-pulldown

#### Blocking beads

200 μl of High-Performance Streptavidin Sepharose beads (GE Healthcare) were washed twice with 1 ml wash buffer-100 (20 mM HEPES-KOH pH 7.9, 100 mM NaCl, 10 mM MgCl_2_ 0,01% (v/v) NP-40, 1 mM DTT) at 4°C and incubated with 1 ml blocking buffer (1 mg/ml BSA, 200 μg/ml Glycogen, 200 μg/ml Yeast tRNA, 0,01% (v/v) NP-40 in wash buffer-100) for 2,5 h at 4°C, washed 3x with wash buffer-300 (20 mM HEPES-KOH pH 7.9, 300 mM NaCl, 10 mM MgCl_2_ 0,01% (v/v) NP-40, 1 mM DTT) and stored in wash buffer-150 (20 mM HEPES-KOH pH 7.9, 150 mM NaCl, 10 mM MgCl_2_ 0,01% (v/v) NP-40, 1 mM DTT) at 4°C until use.

#### Cell lysis and preclearing

##### Pulldown

Approximately 4-8 × 10^9^ cells were washes with cold PBS, centrifuged at 6.000 x g for 10 min and resuspended in lysis buffer-150 (20 mM Tris pH 7.5, 150 mM NaCl, 1,5 mM MgCl_2,_ 2 mM DTT, 2 mM Ribonucleoside Vanadyl Complex (NEB), Roche cOmplete™) to a concentration of 100 μl/10^6^ cells before sonification. The lysate was centrifuged for 15 min at 16.000 x g at 4°C. 10 ml supernatant was precleared by incubation with 200 μl blocked High Performance Streptavidin Sepharose beads for 3 h at 4°C on a rotator wheel.

##### 4xS1m-tagged RNA synthesis

For *in vitro* transcription, the Megascript kits for SP6 and T7 (Invitrogen) were used. In a 40 μl reaction, 2 μg of linearized plasmid (pSP73-4xS1m with different inserts) was used as a template and the reaction was left at 37°C overnight. The next day, 1 μl Turbo DNase was added and the reaction was incubated 15 more min at 37°C, followed by a clean-up with mini quick spin columns (Roche).

##### RNA-pulldown

Each RNA-pulldown was carried out in duplicate to obtain enough protein eluate for LC-MS. Per sample, 50 pmol of *in vitro* transcribed RNA was added to precleared cell lysate (pulldown 1: 2 mg/sample, pulldown 2: 1,5 mg/sample) supplemented with 2,5 μl/ml RNase inhibitor, 0,1 μg/ml tRNA and incubated 1 h at 4°C on a rotator wheel. 10 μl sample from each duplicate was taken for RNA extraction (input). 35 μl blocked beads were added and incubated at 4°C for 1,5 h on a rotator wheel. The beads were settled by centrifugation and 10 μl from each duplicate of the supernatant was taken for RNA extraction (flow-through). The beads were washed 5x with wash buffer and 10 μl bead slurry was taken for RNA extraction. The rest of the beads was incubated with wash buffer supplemented with 50 μg/ml RNase A (Applichem) for 15 min on ice. The eluates were combined in a new Eppendorf tube and supplemented with 1,5 ml ice-cold acetone for protein precipitation and kept at -20°C overnight. The was centrifuged 30 min at 17.000 x g at RT, the pellet washed twice with 80% sterile ethanol (RT), air-dried and resuspended in 36 μl 1x SDS loading buffer. Samples were denatured 5 min at 95°C and kept on ice before proteins were separated on a gel.

### In-gel tryptic digestion and LC-MS/MS analysis

After SDS-PAGE coomassie stained bands were cut out and processed as described previously (Barenz, Inoue et al., 2013). In brief, samples were reduced, alkylated and digested with trypsin. Peptides were extracted from the gel pieces, concentrated in a vacuum centrifuge and dissolved in 15 µl 0.1% TFA. Nanoflow LC-MS2 analysis was performed with an Ultimate 3000 liquid chromatography system coupled to an Orbitrap Elite (Thermo Fisher)with in-house packed analytical column (75 µm x 200 mm, 1.9 µm ReprosilPur-AQ 120 C18 material (Dr. Maisch, Germany). Peptides were separated in a 25 min linear gradient (3-40% B) (Solvent A: 0.1% formic acid / 1% acetonitrile, solvent B: 0.1% formic acid, 89.9% acetonitrile). The mass spectrometer was operated in data-dependent acquisition mode, automatically switching between MS and MS2. MS spectra (m/z 400–1600) were acquired in the Orbitrap at 60,000 (m/z 400) resolution and MS2 spectra were generated for up to 15 precursors with normalized collision energy of 35% in the ion trap. The MS/MS spectra were searched against the uniprotKB Drosophila database (13776 entries) and a contaminants database (MaxQuant) using Proteome Discoverer 2.5 with Sequest (Thermo Fisher Scientific). Carbamidomethyl was set as fixed modification of cysteine and oxidation (methionine), deamidation (asparagines, glutamine) and acetylation (protein N-terminus) as variable modifications. Mass tolerance was set to 5ppm and 0.5 Da for MS and MS/MS, respectively. Only high confident peptides were used and the false discovery rate was set to 0.01.

### RNA-IP

The RNA-IP was performed according to the protocol published in (Tiwari, Zeitler et al., 2019) with GFP-Trap Magentic Particles M-270 (Chromotek) with 50 ovaries collected in 20 µl RNase-free PBS on ice. The PBS was substituted with 40 µl lysis buffer (50 mM Tris pH 7.5, 150 mM NaCl, 5 mM EDTA, 0,5% (v/v) NP-40, 10% (v/v) glycerol, 0,5 mM DTT, Roche cOmplete™, and 5% (v/v) Ribonucleoside Vanadyl Complex (NEB)) and homogenized centrifuged at 15.000 g at 4°C.

Per sample, 25 µl of washed bead slurry was incubated with 500 µl ovary lysate at 4°C for 2 h and washed with lysis buffer and twice with wash buffer (50 mM Tris pH 7.5, 500 mM NaCl, 5 mM EDTA, 0,5% (v/v) NP-40, 10% (v/v) glycerol, 0,5 mM DTT, Roche cOmplete™, and 5% (v/v) Ribonucleoside Vanadyl Complex (NEB)) at 4°C for 10 min. Half of the beads were resuspended in 1x SDS loading buffer for WB analysis and the other half were resuspended in 500 µl Trizol for RNA isolation.

### Western Blot

15 µl per well were loaded onto a 15-well TGX precast gel 4-15% (Biorad) and run 45 min. at 150V. The gel was blotted on a 0.2 µm nitrocellulose membrane (GE healthcare) using the TransBlot Turbo (Biorad) at 1.3A, 25V for 7 min. Membranes were blocked in TBST + 5% milk and incubated with the primary rabbit **α**-YFP antibody (1:5000, (Shieh, Minguez et al., 2015)) in TBST + 5% milk at 4°C overnight. Subsequently, the membrane was washed three times with TBST for 10 min., followed by incubation with the secondary **α**-rabbit HRP-conjugated antibody (1:500, Sigma A0545) for 1 hour at RT. After three more washes with TBST, the membrane was treated with SuperSignal™ West Femto Maximum Sensitivity Substrate (Thermo Scientific) and exposed on the Amersham Imager 600 (GE healthcare).

### Cell culture

Drosophila Schneider 2 (S2) cells were grown in Schneider’s Drosophila medium (Gibco) supplemented with 10% (v/v) Fetal Bovine Serum (PAN) and 200 μg/ml penicillin and streptomycin (Capricorn Scientific) at 25°C. Transfected cells were supplemented with additional selection antibiotics Hygromcin (250 μg/ml) or Puromycin (50 μg/ml) (Sigma Aldrich). For cell growth in suspension, 10 μg/ml Heparin and 0,05% Synperonic (Sigma Aldrich) were added to the medium and cells were incubated in an Erlenmeyer flask while shaking 80 min^-1^.

### Transfections

1,5 × 10^6^ cells/wells were transfected with 5 μg pDNA using Cellfectin II reagent (Gibco) according to the manufacturers protocol.

### Immunofluorescent staining of cells

Immunofluorescence was performed as described (Mathew, Pauleau et al., 2014). Briefly, 3-5 × 10^5^ cells were settled on Polysine Slides (Thermo Fisher), fixed with 4%, washed 3x 5 min with PBS and permeabilised with 0.1% Triton X-100 in PBS for 5 min. Cells were blocked with 4% BSA in PBS for 30 min followed by incubation with the primary antibody diluted in 4% BSA in PBS overnight. Cells were washed 3x with PBS before incubating with the fluorescent secondary antibody diluted 1:500 in 4% BSA in PBS for 1 h protected from light, washed 3x with PBS, counterstained with 1 µg/ml DAPI in PBS, washed again and mounted with Aqua-Poly/Mount (Polysciences). The following antibodies were used: mouse **α**-tubulin (1:1000, Sigma T9026) and rabbit **α**-Cenp-C (Pauleau, Bergner et al., 2019). Secondary fluorescent antibodies were purchased from Thermo Fisher Scientific.

### Drosophila husbandry

The fly stocks were reared and maintained on standard medium at 18°C. Crosses were set up with 2/3 virgins and 1/3 males and kept at 25°C, except for crosses with MTD-Gal4, which were kept at 18 °C. The following fly lines were used: Bloomington 35587 (trip Centagon 1), Bloomington 56998 (trip Centagon 2), Bloomington 33340 (trip Centagon 3), Bloomington 61865 (trip Centagon 4), Bloomington 33997 (trip CG12128), Mat67.15-Gal4 (Bloomington 80361), MTD-Gal4 (Bloomington 31777), Zhr^1^ (origin?), GR1-Gal4 and traffic jam-Gal4 (Veit Riechmann lab), UAS-NLS-GFP (Ingrid Lohmann lab), w^1118^. Endogenously tagged Centagon-GFP flies were purchased from Qidong Fungene Biotechnology. For dissections and fly sorting, a stereomicroscope from Zeiss with external light source was used.

### Generation of transgenic flies

The region of the Centagon 1 gene targeted by the dsRNA of the trip fly line was designed with alternative codons and this altered Centagon 1 gene sequence including its upstream and downstream regions was cloned into the pATTB vector (Ni, Zhou et al., 2011). The plasmid was sent to the fly facility of Cambridge University for injection into vas-int;attp40 embryos and the resulting flies were crossed to obtain a homozygous stock.

### Survival assay

To obtain females with ovary-specific KDs, different UAS-RNAi females were crossed to Mat 67;15-Gal4 males. An equal number of newly hatched F1 flies (30 females and 20 males) were crossed and eggs laid overnight on day tree were collected and transferred to a new grape juice plate with Nipagin (Sigma) and the number of hatched eggs was assessed after 24 h. Eggs were collected 5 consecutive nights to monitor egg laying at different ages.

### Ovary IF

Adult ovaries were dissected in 1x PBS and fixed with 4% PFA in PBS for 20 min. The ovaries were washed 3x 15min in PBST (0,1% Tween in PBS) and permeabilised in 1% Triton-X in PBS for 30 min. After three 15 min washes in PBST, the ovaries were blocked in antibody blocking solution (0,1% (v/v) Tween-20, 0,1% BSA, 10% FBS in PBS) for 2 h, followed by incubation with the primary antibody in PBST at 4°C overnight. Ovaries were washed in PBST 3x 15min, followed by secondary antibody for 2 h. The ovaries were rinsed once and washed twice for 15 min with PBST and counterstained with DAPI (1:000) in PBS. After two more 5 min PBST washes, the ovaries were mounted onto coverslips, with Aqua Polymount mounting medium. The following antibodies were used: rabbit **α**-pMad (1:500, Abcam 52903), rabbit **α**-vasa (1:30, DSHB), mouse **α**-modulo (1:5000, Jaques Pradel lab). Secondary fluorescent antibodies were purchased from Thermo Fisher Scientific.

### Testis IF

Adult testes were dissected in PBS, fixed in 4% formaldehyde for 20 min and blocked in 5% BSA for 1 h at room temperature. Testes were incubated in primary antibodies (in PBST) overnight at 4°C, then washed for 1 h in PBST and incubated with secondary antibodies (in 5% BSA) for 2 h. After washing for 1 h in PBS, testes were mounted on Vectashield (Vector Lab). The following antibodies were used: mouse anti-fasciclin III 7G10 (1:100, DSHB), guinea pig anti traffic-jam (1:5000, Dorothea Godt lab), rabbit anti-vasa (1:250, Santa Cruz d-260). Secondary fuorescent antibodies were purchased from Thermo Fischer Scientific and used 1:500.

### Ovary RNA FISH

For RNA FISH, single molecule Hulu FISH probes from Pixelbio were used according to the manufacturer’s protocol. Dissected ovaries were fixated with 4% PFA for 30 min, washed twice 5 min with PBS and permeabilized in 70% ethanol overnight at 4°C. The next day, the ovaries were washed 2x 10 min with Hulu wash buffer (2x SSC, 2M Urea), before adding 50 μl Hulu hybridization buffer (2x SSC, 2M Urea, 10% dextran sulfate sodium salt, 5x Denhardt’s solution) and 0,5 μl Hulu probe. The ovaries were incubated overnight at 30°C protected from light, then washed 4x 10 min with HULU wash buffer and incubated with 1 μg/μl DAPI in Hulu wash buffer for 5 min and washed 2x 5 min with Hulu wash buffer and mounted with Aqua/Polymount on coverslips. All solutions were RNase-free. For RNase experiments, ovaries were incubated 1h with 20 μg/ml RNase A in PBS or only PBS at 37°C before fixation.

### Microscopic techniques

#### DeltaVision microscope

S2 cells with immunofluorescence staining were imaged with the DeltaVision Core system (Applied Precision) using the Olympus UPlanSApo 100x (n.a. 1.4). Z-slices were 0,2 μm. Deconvolution was performed with the Applied Precisions softWoRx 3.7.1 suite with the following settings: Ratio (conservative), 10 cycles. Images used as examples in figures were adjusted for brightness and contrast in FIJI (ImageJ).

#### Confocal microscopy

For Drosophila tissues, either the Leica TCS SP5II confocal microscope with the HCX Plan APO 40x/1.30 Oil Cs objective was used or the Leica TSC SP8 confocal microscope with the PLAN APO 20x, multi-immersion (NA 0.75) and PLAN APO 63x, glycerol immersion (NA 1.3) objectives. 2 μm z-slices were imaged for whole ovaries and egg chamber and 0,5 μm z-slices were used for the imaging of germaria. Images used as examples in figures were adjusted for brightness and contrast in FIJI (ImageJ).

#### Quantifications

Microscopic images were analysed in FIJI (ImageJ). For intensity measurements, background was removed with the rolling ball tool and a z-projection of the z-slices with the signal of interest was made. Then, the region of interest was selected and the Raw Int Den was measured.

##### sat III signal in egg chambers

The fragmentation of nurse cell chromatin was assessed by eye and egg chambers with comparable sizes were analysed.

##### pMad levels in MTD-Gal4-induced KD ovaries

GSCs were selected one by one for intensity measurements. Cystoblasts with pMad signal were counted and all pMad-expressing cystoblasts were selected together for one intensity measurement. The obtained value was divided by the number of cells counted in the selected area.

## Acknowledgements

We thank the Bloomington stock center for providing transgenic fly stocks, the TRiP at Harvard Medical School (NIH/NIGMS R01-GM084947) for providing transgenic RNAi fly stocks and V. Riechmann for gifting germ line-specific Gal4-driver lines. We thank J. Pradel lab for anti-modulo antibodies, B. Bukau lab for anti-YFP antibodies, and the D. Godt lab for the anti-traffic jam antibodies. The anti-vasa and anti-fascilin III antibodies developed by Spradling/Williams and Goodman, respectively, were obtained from the Developmental Studies Hybridoma bank, created by the NICHD of the NIH and maintained at the university of Iowa, Department of Biology, Iowa City, IA 52242. We thank G. Stoecklin for the S1m aptamer constructs, S. Diederichs’ lab for their RNA-pulldown protocol, the ZMBH Imaging Facility for microscopy assistance and the ZMBH proteomics facility for the Mass spec analysis. Many thanks to V. Riechmann, G. Stoecklin, A. Maizel and the Erhardt and Lohmann laboratory members for insights and discussions, and S. Corless and A.L. Pauleau for critical comments on the manuscript. We acknowledge funding from the Deutsche Forschungsgemeinschaft (DFG) through the grant EXC81 (CellNetworks), SFB1036, the European Research Council through the grant ERC-CoG-682496 (cenRNA) to SE, the European Commission Erasmus+ exchange program for supporting LAK. SLH is an alumna and AMS a member of the HBIGS graduate school program.

## Author contributions

Saskia Höcker: Conceptualization, Investigation, Resources, Data curation, Software, Formal analysis, Supervision, Validation, Visualization, Methodology, Writing—original draft, editing; Izlem SuAkan: Investigation, Visualization, Methodology; Alexander Simon: Investigation, Formal analysis; Lili Kenez: Investigation; Kirem Yildirim: Investigation, Visualization, Methodology. Ingrid Lohmann: Supervision, editing; Sylvia Erhardt: Conceptualization, Supervision, Funding acquisition, Project administration, Investigation, Writing—original draft, review and editing

## Competing interests

No competing interests declared

## Supplemental Information Figure Legends

**Supplemental Figure S1.**
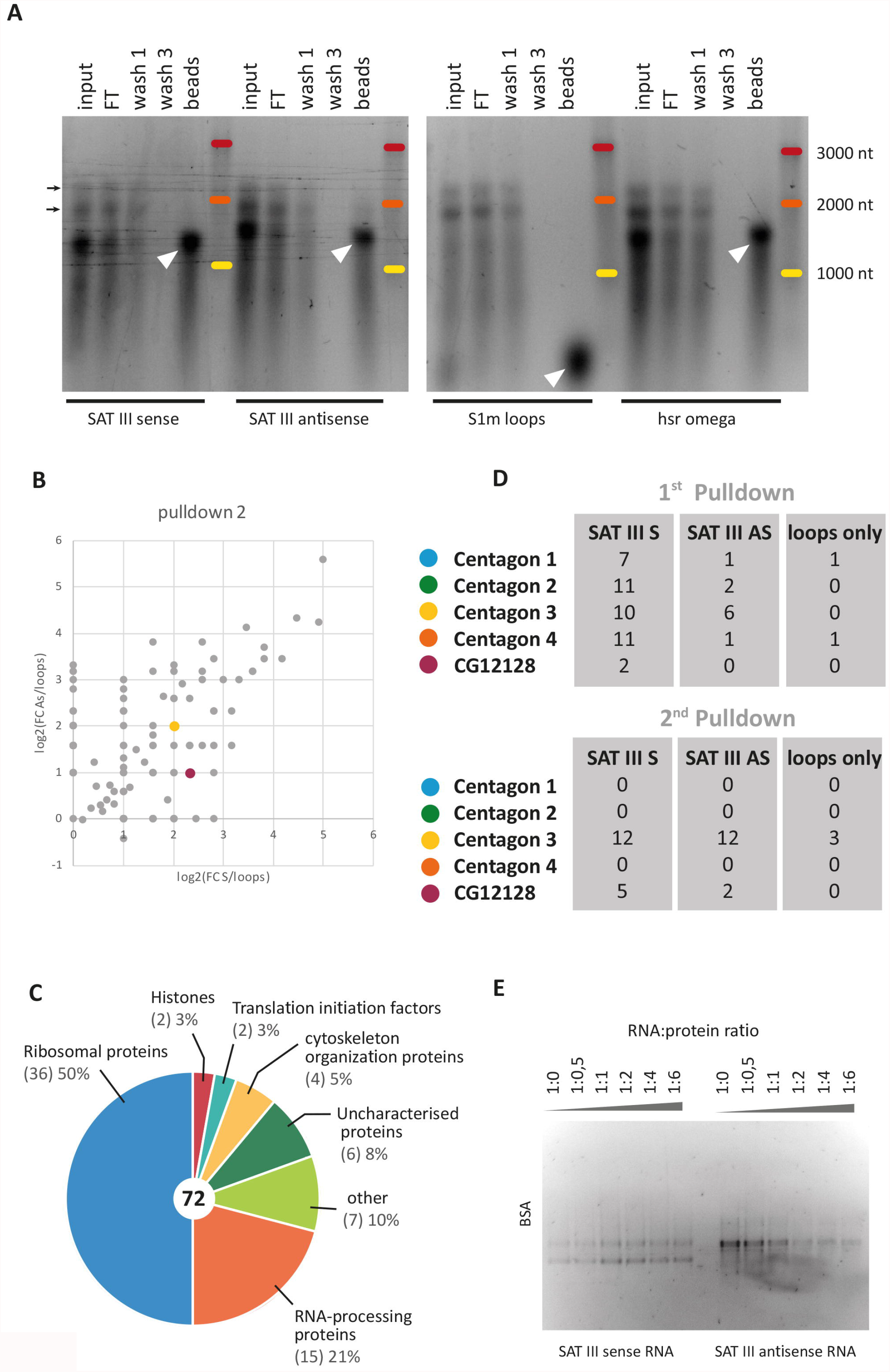
**A**. Denaturing gel with RNA of the different RNA-pulldown steps. The S1m-loop tagged RNA attached to the beads is indicated by a white triangle. The black arrows indicate rRNA bands in the input and flow-through samples which derive from the S2 cell lysate. **B**. Enrichment of proteins in the *SAT III sense* and *antisense* RNA pulldown compared to 4xS1m loop RNA pulldown in the second RNA-pulldown experiment. Each dot represents a protein and its fold change (FC) in the *SAT III sense* RNA pulldown (x-axis) and *SAT III antisense* RNA pulldown (y-axis) compared to the control on a logarithmic scale. Coloured dots represent members of the Centagon complex or the CG12128 control protein. **C**. Overview of the protein classes enriched in both *SAT III sense* RNA pulldowns. The number of identified proteins in each class is indicated in brackets. **D**. Overview of the peptide counts of the Centagon proteins in the MS analysis of each RNA pulldown experiment. **E**. EMSA of *SAT III sense* and *antisense* RNA with BSA. *In vitro* transcribed RNA was incubated with BSA in the indicated molar ratios.

**Supplemental Figure S2.**
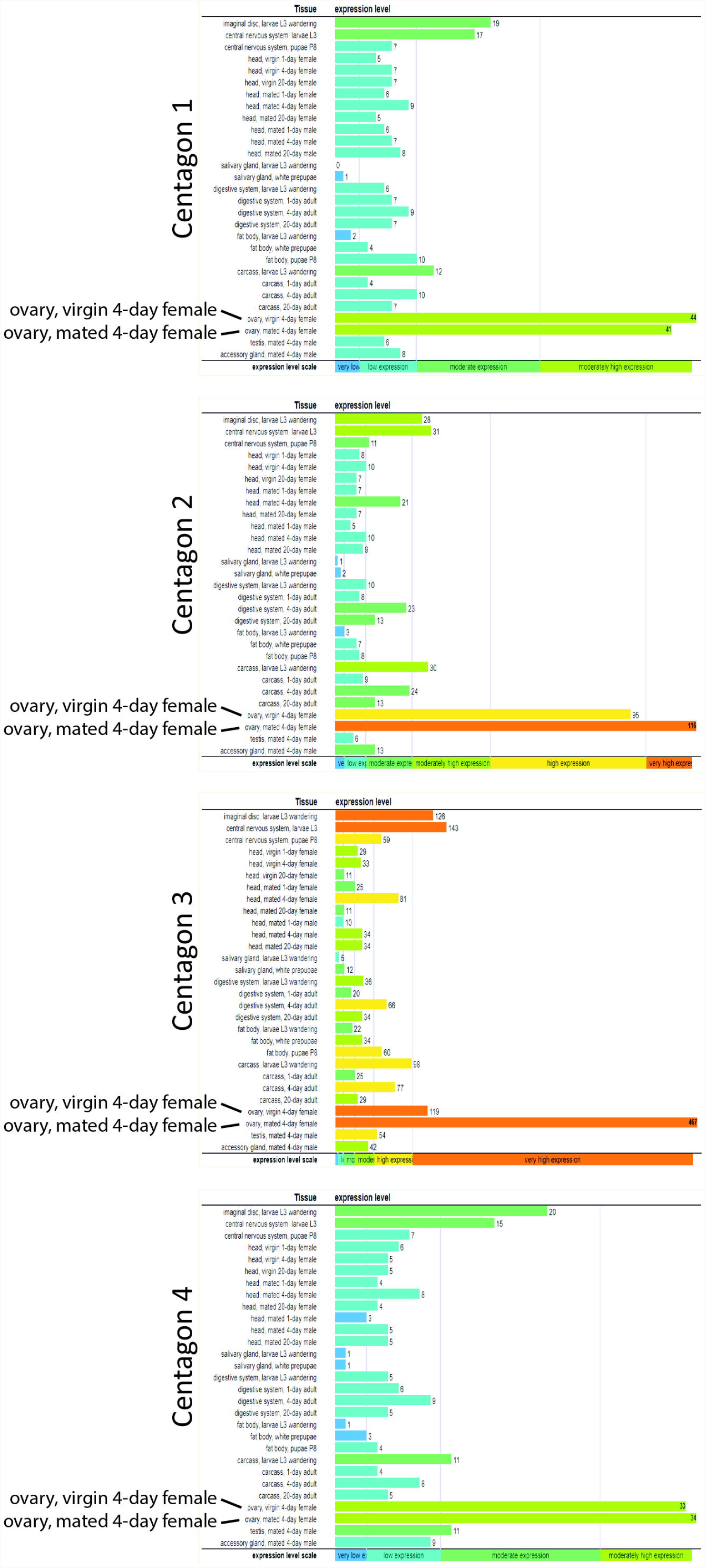
Expression data of the Centagon members per tissue according to www.flybase.org.

**Supplemental Figure S3.**
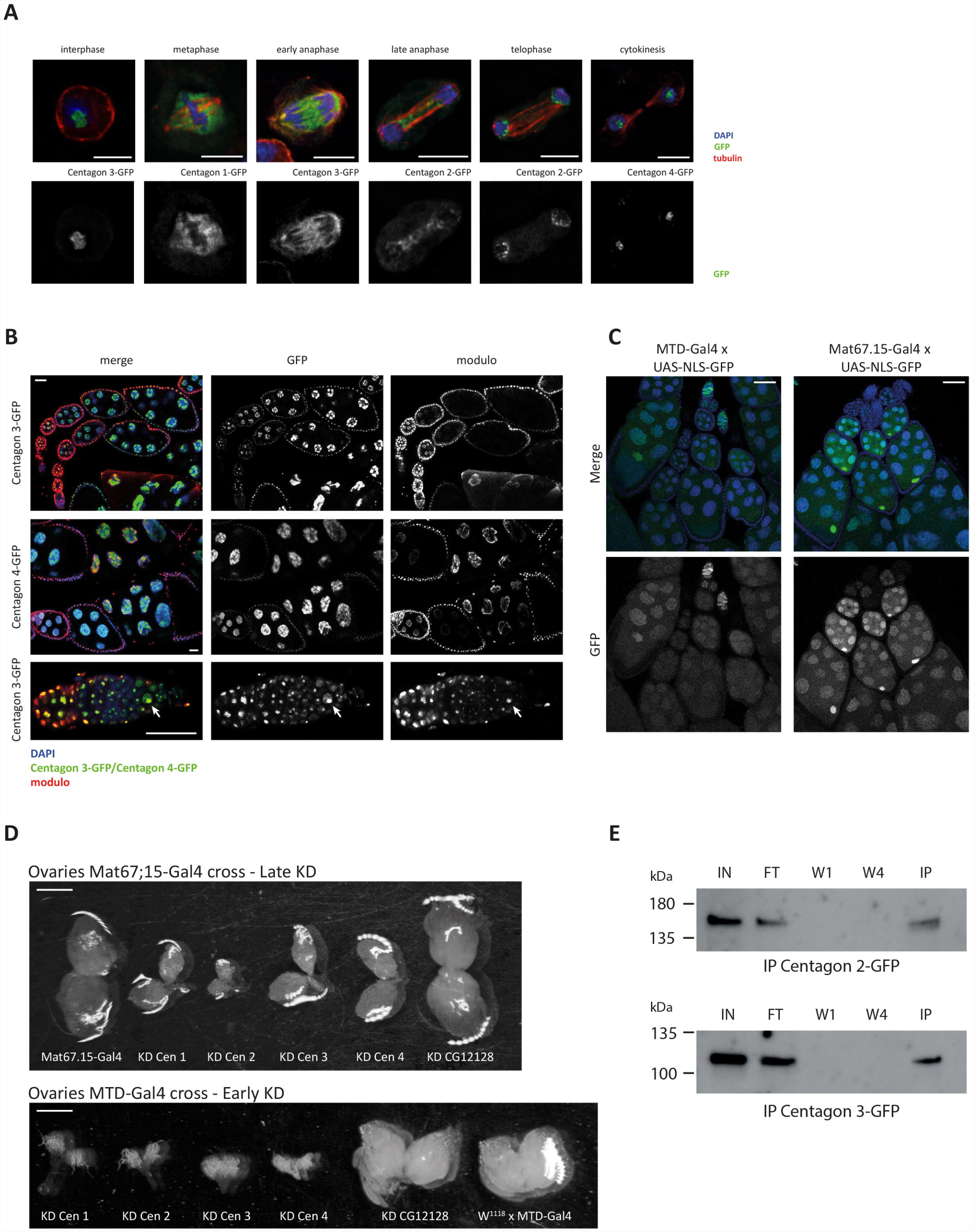
**A**. Exogenously GFP-tagged Centagon proteins imaged throughout the cell cycle in S2 cells with the different Centagon proteins (as indicated) in green, anti-tubulin in red and DNA in blue. Scale bar 5 μm. **B**. Ovarioles (top panels) and germarium (bottom panel) of flies with endogenous GFP-tagged Centagon proteins (green), co-immunostained for the nucleolar protein Modulo (red) and DAPI (blue). Scale bar 20 μm. **C**. F1 ovaries from the parent cross: UAS-NLS-GFP x Mat67.15-Gal4 or MTD-Gal4, stained with DAPI (blue). The GFP-tagged Nuclear Localization Sequence (NLS) (green) shows at what point in ovary development the driver line starts Gal4 expression in germ cells: The MTD driver in the germarium, the Mat67.15 driver from early egg chambers on. Scale bar 50 μm. **D**. Ovaries dissected from MTD-or Mat67.15-induced Centagon KD females. Scale bar 0,5 mm. **E**. Western Blot of the different steps in the RNA-IP shown in Fig. 3F-G. From left to right: Input, flowthrough, wash 1, wash 4 and IP. A YFP-antibody was used to detect the GFP-tagged Centagon proteins.

**Supplemental Figure S4.**
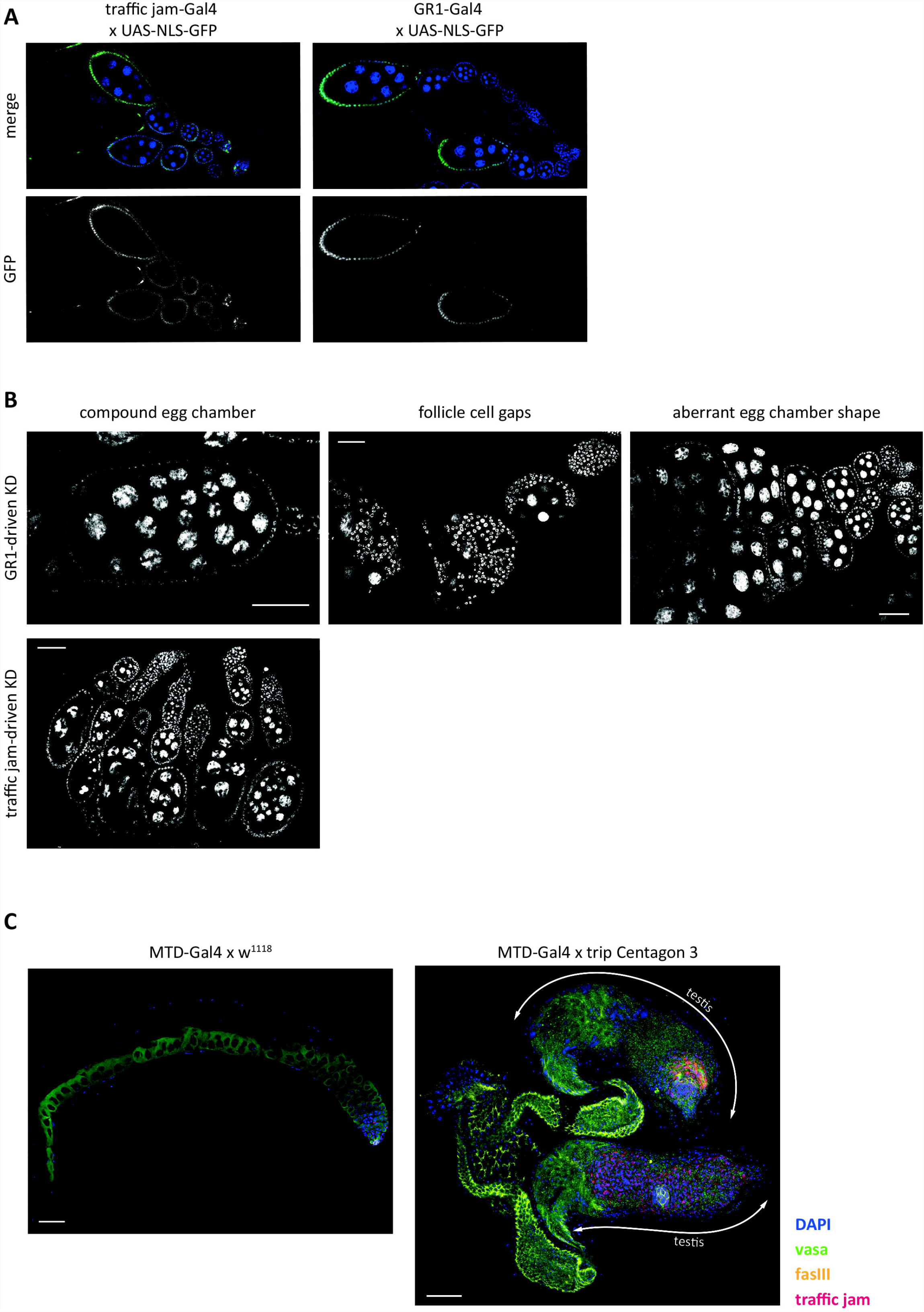
**A**. UAS-NLS-GFP flies were either crossed with traffic jam-Gal4 or GR1-Gal4, and F1 ovaries were stained with DAPI (blue). The GFP-tagged Nuclear Localization Sequence (NLS) (green) appears at the developmental stage when the driver line starts expressing Gal4 in somatic cells: The traffic jam driver in the germarium, the GR1 driver in the follicle cells of later egg chambers. Scale bar 50 μm. **B**. Examples of DAPI-stained ovaries dissected from GR1-induced Centagon KD females with the indicated phenotypes. Scale bar 50 μm. **C**. Control (left) and Centagon 3 KD testis (right), induced with the MTD-Gal4 driver. Immunofluorescent staining of the hub (anti-fas III, yellow), the germ cells (anti-vasa, green) and cyst cells (traffic jam, red), co-stained with DAPI (blue). Scale bar 50 μm.

**Supplemental Figure S5.**
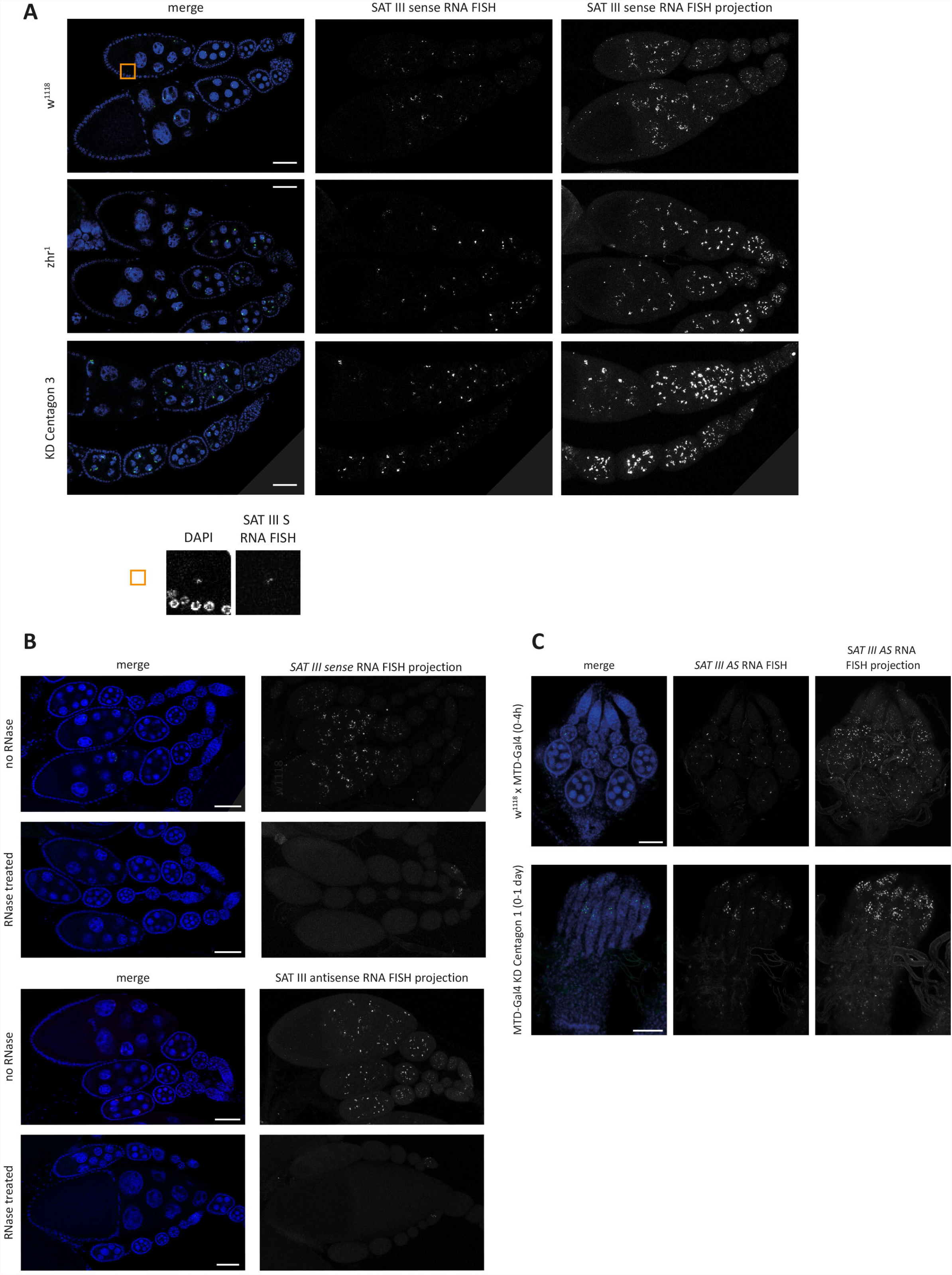
**A**. *SAT III sense* RNA FISH (green) in w^1118^, Zhr^1^ and Mat67.15-Gal4-induced Cen3 KD ovaries, stained with DAPI (blue). *SAT III* RNA signals are depicted in one z-slice (middle) and as z-stack projection (right). The orange square enlarged on top shows the oocyte nucleus. For easier orientation, some images were rotated on top of a grey background. Scale bar 50 μm. **B**. *SAT III sense* and *antisense* RNA FISH in w^1118^ ovaries with or without RNase A digestion, stained with DAPI. The *SAT III* RNA FISH signal is shown as a z-stack projection. Scale bar 50 μm. **C**. *SAT III antisense* RNA FISH on newly hatched control (0-4 h old) and Centagon 1 KD (0-1 day old) ovaries. SAT III RNA signals are depicted in one z-slice (middle) and as z-stack projection (right). Scale bar 50 μm.

## Supplemental tables

### Mass Spectromerty results of the *SAT III* RNA pulldowns

**Table 1: pulldown 1**

Peptide counts in *SAT III sense* (S), *antisense* (AS) and ‘*loops only’* (L) RNA pulldown. log2(FC) = log2 values of the fold change of S/L, only proteins with a log2(FC) of 1 or higher are listed.

**Table 2: pulldown 2**

Peptide counts in *SAT III sense* (S), *antisense* (AS) and ‘*loops only’* (L) RNA pulldown. log2(FC) = log2 values of the fold change of S/L, only proteins with a log2(FC) of 1 or higher are listed.

## Notes

### Competing Interest Statement

The authors have declared no competing interest.

