## Supplemental Table S1 for "A centromeric RNA-associated protein complex affects germ line development in *Drosophila melanogaster*"

### Table S1: Sat III RNA pulldown 1 - MS data

S = peptide counts in sat III sense RNA pulldown
AS = peptide counts in sat III sense RNA pulldown
L = peptide counts in loops only RNA pulldown
log2(FC) = log2 values of the fold change of S/L

Cutoff: only proteins with a log2(FC) of 1 or higher a listed

| CG number | protein ID | S | AS | L | log2(FC) |
|---|---|---|---|---|---|
| CG9684 | CG9684 | 30 | 16 | 0 | 4,9 |
| CG13900 | CG13900-PA, isoform A | 25 | 8 | 1 | 4,6 |
| CG5787 | CG5787, isoform A | 20 | 14 | 0 | 4,3 |
| CG2691 | RRP12-like protein | 19 | 2 | 1 | 4,2 |
| CG8710 | Isoform of A1Z7A8, Coilin, isoform E | 17 | 0 | 0 | 4,1 |
| CG1828 | FACT complex subunit spt16 | 16 | 3 | 0 | 4 |
| CG3605 | CG3605, isoform A | 15 | 2 | 0 | 3,9 |
| CG5931 | Putative U5 small nuclear ribonucleoprotein 200 kDa helicase | 15 | 0 | 1 | 3,9 |
| CG5720 | BcDNA.LD27873 | 15 | 8 | 1 | 3,9 |
| CG6905 | Cell division cycle 5 ortholog, isoform A | 13 | 3 | 1 | 3,7 |
| CG2807 | CG2807, isoform A | 25 | 6 | 2 | 3,6 |
| CG17838 | Isoform of A0A0B4KHI4, Syncrip, isoform O | 12 | 0 | 0 | 3,6 |
| CG12785 | Nucleolar protein 6 | 12 | 0 | 0 | 3,6 |
| CG6189 | FI21448p1 | 12 | 0 | 1 | 3,6 |
| CG8545 | CG8545 | 11 | 2 | 0 | 3,5 |
| CG12819 | Protein slender lobes | 11 | 4 | 0 | 3,5 |
| CG12085 | Poly(U)-binding-splicing factor half pint | 11 | 3 | 0 | 3,5 |
| CG32344 | CG32344 | 11 | 1 | 1 | 3,5 |
| CG13096 | Isoform of Q9VLK2, CG13096, isoform B | 10 | 6 | 0 | 3,3 |
| CG8877 | Pre-mRNA processing factor 8 | 10 | 1 | 1 | 3,3 |
| CG6711 | Isoform of Q24325, LD23043p | 9 | 0 | 0 | 3,2 |
| CG2009 | CG2009-PA | 9 | 2 | 0 | 3,2 |
| CG6701 | CG6701, isoform B | 9 | 3 | 0 | 3,2 |
| CG17603 | Isoform of P51123, TBP-associated factor 1, isoform F | 9 | 4 | 0 | 3,2 |
| CG5786 | Protein Peter pan | 9 | 1 | 1 | 3,2 |
| CG1091 | Isoform of A0A0B4KGN4, CG1091, isoform B | 9 | 1 | 1 | 3,2 |
| CG9226 | Isoform of Q6NL34, WD repeat domain 79 homolog, isoform B | 8 | 0 | 0 | 3 |
| CG4817 | FACT complex subunit Ssrp1 | 8 | 0 | 0 | 3 |
| CG7752 | CG7752-PA | 8 | 3 | 1 | 3 |
| CG11207 | CG11207-PA | 8 | 1 | 1 | 3 |
| CG7757 | Precursor RNA processing 3, isoform A | 8 | 0 | 1 | 3 |
| CG8801 | Isoform of Q9V411, Nucleolar GTP-binding protein 1 | 7 | 2 | 0 | 2,8 |
| CG10269 | D19A | 7 | 0 | 1 | 2,8 |
| CG8174 | SRPK, isoform D | 7 | 2 | 1 | 2,8 |
| CG1234 | Nucleolar complex protein 3 homolog | 7 | 1 | 1 | 2,8 |
| CG6671 | Argonaute-1, isoform A | 14 | 0 | 2 | 2,8 |
| CG7704 | Transcription initiation factor TFIID subunit 5 | 7 | 1 | 1 | 2,8 |
| CG10042 | MBD-R2 | 7 | 5 | 1 | 2,8 |
| CG9750 | RuvB-like helicase 2 | 7 | 0 | 1 | 2,8 |
| CG6995 | Scaffold attachment factor B, isoform B | 6 | 0 | 0 | 2,6 |
| CG1785 | Uncharacterized protein CG1785 | 6 | 2 | 1 | 2,6 |
| CG31739 | CG31739, isoform A | 6 | 1 | 1 | 2,6 |
| CG1685 | Protein penguin | 17 | 2 | 3 | 2,5 |
| CG10600 | CG10600, isoform B | 5 | 1 | 0 | 2,3 |
| CG7831 | Protein claret segregational | 15 | 11 | 3 | 2,3 |
| CG6937 | CG6937 | 5 | 1 | 1 | 2,3 |
| CG11522 | RE08669p | 5 | 3 | 1 | 2,3 |
| CG14938 | CG14938-PA, isoform A | 5 | 0 | 1 | 2,3 |
| CG9916 | Peptidyl-prolyl cis-trans isomerase | 5 | 0 | 1 | 2,3 |
| CG7977 | FI01658p | 5 | 8 | 1 | 2,3 |
| CG1796 | LD24662p | 5 | 1 | 1 | 2,3 |

| | | | | | |
|---|---|---|---|---|---|
| CG2260 | CG2260 | 4 | 0 | 0 | 2 |
| CG10333 | CG10333 | 4 | 1 | 0 | 2 |
| CG16725 | Survival motor neuron protein | 4 | 0 | 0 | 2 |
| CG10712 | Chromator, isoform A | 4 | 1 | 0 | 2 |
| CG4951 | Uncharacterized protein CG4951 | 4 | 2 | 0 | 2 |
| CG1258 | Kinesin-like protein | 4 | 0 | 0 | 2 |
| CG6605 | Isoform of P16568, Bicaudal D, isoform D | 4 | 0 | 0 | 2 |
| CG8611 | Probable ATP-dependent RNA helicase CG8611 | 4 | 1 | 0 | 2 |
| CG1832 | Isoform of Q9I7K0, CG1832-PA, isoform A | 4 | 0 | 0 | 2 |
| CG10270 | D19B | 4 | 0 | 0 | 2 |
| CG8332 | Isoform of Q7JZW2, Ribosomal protein S15, isoform B | 4 | 1 | 1 | 2 |
| CG4863 | 60S ribosomal protein L3 | 12 | 7 | 3 | 2 |
| CG9630 | Probable ATP-dependent RNA helicase DDX55 homolog | 4 | 0 | 1 | 2 |
| CG9888 | rRNA 2’-O-methyltransferase fibrillarin | 8 | 0 | 2 | 2 |
| CG1866 | Moca-cyp, isoform A | 4 | 0 | 1 | 2 |
| CG17136 | RNA-binding protein 1 | 4 | 0 | 1 | 2 |
| CG9755 | Maternal protein pumilio | 7 | 1 | 2 | 1,8 |
| CG3231 | Something that sticks like glue, isoform A | 7 | 2 | 2 | 1,8 |
| CG32211 | Transcription initiation factor TFIID subunit 6 | 10 | 5 | 3 | 1,7 |
| CG6322 | SD09427p | 10 | 1 | 3 | 1,7 |
| CG7728 | CG7728-PA | 3 | 0 | 0 | 1,6 |
| CG8817 | AF4/FMR2 family member 4 | 3 | 2 | 0 | 1,6 |
| CG4616 | FLASH ortholog, isoform A | 3 | 0 | 0 | 1,6 |
| CG6546 | Brahma associated protein 55kD | 3 | 0 | 0 | 1,6 |
| CG7518 | Isoform of Q9VG05, CG7518, isoform F | 3 | 2 | 0 | 1,6 |
| CG8233 | Isoform of A0A0B4KF25, Reduction in Cnn dots 1, isoform H | 3 | 2 | 0 | 1,6 |
| CG6988 | Isoform of P54399, Protein disulfide-isomerase | 3 | 0 | 0 | 1,6 |
| CG31012 | Isoform of A0A0B4KI34, CIN85 and CD2AP orthologue, isoform F | 3 | 1 | 0 | 1,6 |
| CG32763 | CG32763-PA | 3 | 1 | 0 | 1,6 |
| CG1542 | Probable rRNA-processing protein EBP2 homolog | 3 | 0 | 0 | 1,6 |
| CG4699 | Isoform of E2QD16, Non-specific lethal 1, isoform D | 3 | 1 | 0 | 1,6 |
| CG18273 | CG18273 | 3 | 0 | 0 | 1,6 |
| CG17611 | Eukaryotic translation initiation factor 6 | 3 | 1 | 0 | 1,6 |
| CG17064 | Guanylate kinase-associated protein mars | 3 | 2 | 0 | 1,6 |
| CG30149 | Protein rigor mortis | 3 | 0 | 0 | 1,6 |
| CG13345 | FI24033p1 | 3 | 0 | 0 | 1,6 |
| CG16940 | CG16940-PC, isoform C | 9 | 5 | 3 | 1,6 |
| CG11949 | Protein 4.1 homolog | 3 | 1 | 1 | 1,6 |
| CG10354 | 5’-3’ exoribonuclease 2 homolog | 3 | 0 | 1 | 1,6 |
| CG33106 | Isoform of Q9VCA8, Multiple ankyrin repeats single KH domain, isoform D | 3 | 1 | 1 | 1,6 |
| CG10922 | 40S ribosomal protein S10b | 6 | 0 | 2 | 1,6 |
| CG11271 | 40S ribosomal protein S12 | 3 | 0 | 1 | 1,6 |
| CG2986 | 40S ribosomal protein S21 | 3 | 0 | 1 | 1,6 |
| CG3203 | 60S ribosomal protein L17 | 6 | 4 | 2 | 1,6 |
| CG7035 | Nuclear cap-binding protein subunit 1 | 3 | 0 | 1 | 1,6 |
| CG3613 | Isoform of Q9W255, Quaking related 58E-1, isoform D | 3 | 1 | 1 | 1,6 |
| CG2998 | Isoform of Q9W334, Ribosomal protein S28b, isoform B | 3 | 0 | 1 | 1,6 |
| CG6510 | 60S ribosomal protein L18a | 6 | 3 | 2 | 1,6 |
| CG9373 | FI21236p1 | 20 | 11 | 7 | 1,5 |
| CG16901 | RNA-binding protein squid | 17 | 5 | 6 | 1,5 |
| CG8108 | CG8108, isoform A | 8 | 2 | 3 | 1,4 |
| CG7434 | 60S ribosomal protein L22 | 8 | 17 | 3 | 1,4 |
| CG15792 | Myosin heavy chain, non-muscle | 8 | 0 | 3 | 1,4 |
| CG5208 | Protein associated with topo II related-1, isoform A | 8 | 1 | 3 | 1,4 |
| CG7439 | Protein argonaute-2 | 28 | 9 | 11 | 1,3 |
| CG3751 | 40S ribosomal protein S24 | 5 | 0 | 2 | 1,3 |
| CG9715 | CG9715 | 5 | 2 | 2 | 1,3 |
| CG6474 | Transcription initiation factor TFIID subunit 9 | 5 | 2 | 2 | 1,3 |
| CG3314 | Isoform of P46223, Ribosomal protein L7A, isoform E | 5 | 3 | 2 | 1,3 |
| | Histone H4 | 5 | 0 | 2 | 1,3 |
| CG4806 | CG4806 | 17 | 5 | 7 | 1,3 |

| | | | | | |
|---|---|---|---|---|---|
| CG5519 | BcDNA.LD02793 | 19 | 5 | 8 | 1,2 |
| CG18811 | Caprin homolog | 7 | 0 | 3 | 1,2 |
| CG17521 | 60S ribosomal protein L10 | 7 | 7 | 3 | 1,2 |
| CG7993 | Ribosome production factor 2 homolog | 7 | 2 | 3 | 1,2 |
| CG12505 | Activity-regulated cytoskeleton associated protein 1 | 7 | 0 | 3 | 1,2 |
| CG4003 | RuvB-like helicase 1 | 9 | 0 | 4 | 1,2 |
| CG4464 | 40S ribosomal protein S19a | 9 | 3 | 4 | 1,2 |
| CG5920 | Isoform of P31009, Ribosomal protein S2, isoform B | 11 | 4 | 5 | 1,1 |
| CG1691 | Isoform of M9NF14, IGF-II mRNA-binding protein, isoform K | 19 | 3 | 9 | 1,1 |
| | Histone H3 | 2 | 0 | 0 | 1 |
| CG1965 | CG1965, isoform A | 2 | 0 | 0 | 1 |
| CG1559 | Isoform of Q9VYS3, Upf1, isoform B | 2 | 1 | 0 | 1 |
| CG7006 | 60S ribosome subunit biogenesis protein NIP7 homolog | 2 | 0 | 0 | 1 |
| CG9775 | CG9775, isoform A | 2 | 0 | 0 | 1 |
| CG6987 | LD40489p | 2 | 0 | 0 | 1 |
| CG4918 | 60S acidic ribosomal protein P2 | 2 | 1 | 0 | 1 |
| CG10805 | HEAT repeat-containing protein 1 homolog | 2 | 0 | 0 | 1 |
| CG11563 | CG11563 | 2 | 0 | 0 | 1 |
| CG31938 | CG31938-PA | 2 | 0 | 0 | 1 |
| | Histone H1 | 2 | 3 | 0 | 1 |
| CG4364 | Pescadillo homolog | 2 | 0 | 0 | 1 |
| CG5589 | CG5589 | 2 | 0 | 0 | 1 |
| CG3163 | CG3163 | 2 | 0 | 0 | 1 |
| CG12499 | CG12499 | 2 | 0 | 0 | 1 |
| CG32435 | CLIP-associating protein | 2 | 0 | 0 | 1 |
| CG13298 | Splicing factor 3B subunit 6-like protein | 2 | 0 | 0 | 1 |
| CG10473 | Acinus, isoform A | 2 | 0 | 0 | 1 |
| CG31368 | CG31368, isoform D | 2 | 1 | 0 | 1 |
| CG8264 | Isoform of P39736, Bx42, isoform B | 2 | 1 | 0 | 1 |
| CG4602 | LD29830p | 2 | 0 | 0 | 1 |
| CG9246 | Nucleolar complex protein 2 homolog | 2 | 0 | 0 | 1 |
| CG2670 | Isoform of Q9VHY5, TBP-associated factor 7, isoform B | 2 | 0 | 0 | 1 |
| CG4211 | Protein no-on-transient A | 2 | 0 | 0 | 1 |
| CG9143 | CG9143 | 2 | 0 | 0 | 1 |
| CG3780 | RE50839p | 2 | 0 | 0 | 1 |
| CG10923 | Kinesin-like protein | 2 | 0 | 0 | 1 |
| CG6751 | No child left behind | 2 | 0 | 0 | 1 |
| CG6197 | FI18620p1 | 2 | 0 | 0 | 1 |
| CG12128 | CG12128, isoform A | 2 | 0 | 0 | 1 |
| CG1101 | LD24793p | 2 | 0 | 0 | 1 |
| CG10341 | Isoform of Q9VJ62, CG10341, isoform C | 2 | 0 | 0 | 1 |
| CG9213 | Isoform of Q9VXT5, CG9213, isoform B | 2 | 0 | 0 | 1 |
| CG4051 | Egalitarian, isoform B | 2 | 1 | 0 | 1 |
| CG4913 | ENL/AF9-related, isoform B | 2 | 2 | 0 | 1 |
| CG9998 | Isoform of Q24562, U2 small nuclear riboprotein auxiliary factor 50, isoform B | 2 | 0 | 0 | 1 |
| CG8103 | Chromodomain-helicase-DNA-binding protein Mi-2 homolog | 2 | 0 | 0 | 1 |
| CG7622 | 60S ribosomal protein L36 | 2 | 0 | 0 | 1 |
| CG17420 | Isoform of O17445, Ribosomal protein L15 | 2 | 1 | 1 | 1 |
| CG7185 | Cleavage and polyadenylation specificity factor subunit CG7185 | 2 | 0 | 1 | 1 |
| CG9641 | CG9641, isoform A | 2 | 0 | 1 | 1 |
| CG11583 | Ribosome biogenesis protein BRX1 homolog | 2 | 1 | 1 | 1 |
| CG9946 | Isoform of P41374, Eukaryotic translation initiation factor 2alpha, isoform B | 2 | 1 | 1 | 1 |
| CG16753 | CG16753-PA | 2 | 0 | 1 | 1 |
| CG2720 | CG2720-PA | 2 | 0 | 1 | 1 |
| CG4709 | Zinc finger CCCH-type with G patch domain-containing protein | 2 | 0 | 1 | 1 |
| CG13425 | Bancal, isoform C | 2 | 0 | 1 | 1 |
| CG18572 | Isoform of P05990, Rudimentary, isoform D | 2 | 0 | 1 | 1 |
| CG5726 | CG5726 O | 2 | 0 | 1 | 1 |
| CG7843 | Isoform of Q9V9K7, Ars2, isoform E | 2 | 0 | 1 | 1 |
| CG11696 | CG11696 O | 2 | 1 | 1 | 1 |
| CG6692 | Isoform of Q95029, Cysteine proteinase-1, isoform D | 2 | 0 | 1 | 1 |

| | | | | | |
|---|---|---|---|---|---|
| CG1622 | CG1622, isoform A | 2 | 0 | 1 | 1 |
| CG10305 | Isoform of P13008, Ribosomal protein S26, isoform D | 2 | 0 | 1 | 1 |
| CG5729 | Dgp-1, isoform A | 2 | 0 | 1 | 1 |
| CG33505 | LD17611p | 4 | 0 | 2 | 1 |
| CG10851 | Isoform of P26686, B52, isoform O | 6 | 1 | 3 | 1 |
| CG2746 | 60S ribosomal protein L19 | 6 | 4 | 3 | 1 |
| CG4849 | CG4849 | 10 | 0 | 5 | 1 |
| CG3661 | 60S ribosomal protein L23 | 4 | 2 | 2 | 1 |
| CG3922 | 40S ribosomal protein S17 | 8 | 2 | 4 | 1 |
| CG13849 | FI04781p | 10 | 0 | 5 | 1 |
| CG10206 | CG10206-PA | 12 | 0 | 6 | 1 |
| CG3195 | RE28824p | 4 | 6 | 2 | 1 |
| CG14224 | LD38919p | 6 | 5 | 3 | 1 |
| CG12598 | Double-stranded RNA-specific editase Adar | 6 | 1 | 3 | 1 |
| CG3735 | CG3735, isoform A | 8 | 2 | 4 | 1 |
| CG5258 | Isoform of Q9V3U2, NHP2, isoform B | 4 | 0 | 2 | 1 |
