## Supplemental Table S2 for "A centromeric RNA-associated protein complex affects germ line development in *Drosophila melanogaster*"

### Table S2: Sat III RNA pulldown 2 - MS data

S = peptide counts in sat III sense RNA pulldown
AS = peptide counts in sat III sense RNA pulldown
L = peptide counts in loops only RNA pulldown
log2(FC) = log2 values of the fold change of S/L

Cutoff: only proteins with a log2(FC) of 1 or higher a listed

| CG number | protein ID | S | AS | L | log2(FC) |
|---|---|---|---|---|---|
| CG5394 | Bifunctional glutamate/proline--tRNA ligase | 32 | 48 | 0 | 5 |
| CG15100 | Methionyl-tRNA synthetase | 30 | 19 | 0 | 4,9 |
| CG11471 | Isoleucyl-tRNA synthetase, isoform A | 22 | 20 | 0 | 4,5 |
| CG5728 | CG5728 | 18 | 11 | 0 | 4,2 |
| CG3821 | Aspartyl-tRNA synthetase, isoform A | 14 | 11 | 0 | 3,8 |
| CG3178 | Recombination repair protein 1 | 28 | 26 | 2 | 3,8 |
| CG12141 | Lysine--tRNA ligase | 12 | 9 | 0 | 3,6 |
| CG9020 | Probable arginine--tRNA ligase, cytoplasmic | 12 | 9 | 0 | 3,6 |
| CG10302 | Bicoid mRNA stability factor | 22 | 35 | 2 | 3,5 |
| CG15792 | Zipper, isoform C | 10 | 8 | 1 | 3,3 |
| CG33123 | SD07726p | 9 | 5 | 0 | 3,2 |
| CG10279 | LP18603p | 9 | 3 | 1 | 3,2 |
| CG7070 | Pyruvate kinase | 8 | 8 | 1 | 3 |
| CG10206 | CG10206 protein | 7 | 2 | 0 | 2,8 |
| CG8470 | GH05406p | 7 | 5 | 0 | 2,8 |
| CG13900 | RE01065p | 7 | 5 | 1 | 2,8 |
| CG7439 | Protein argonaute-2 | 7 | 0 | 1 | 2,8 |
| CG7490 | 60S acidic ribosomal protein P0 | 7 | 11 | 0 | 2,8 |
| CG8258 | CG8258 | 7 | 3 | 0 | 2,8 |
| CG5599 | Dihydrolipoamide acetyltransferase component of pyruvate dehydrogenase complex | 6 | 9 | 0 | 2,6 |
| CG30122 | GM15464p | 6 | 3 | 1 | 2,6 |
| CG5502 | 60S ribosomal protein L4 | 6 | 14 | 0 | 2,6 |
| CG6701 | CG6701, isoform B | 6 | 8 | 0 | 2,6 |
| CG1691 | IGF-II mRNA-binding protein, isoform D | 11 | 2 | 2 | 2,5 |
| CG12128 | Uncharacterized protein, isoform A | 5 | 2 | 0 | 2,3 |
| CG6203 | Fmr1, isoform G | 5 | 3 | 1 | 2,3 |
| CG17369 | V-type proton ATPase subunit B | 5 | 6 | 1 | 2,3 |
| CG5519 | BcDNA.LD02793 | 5 | 3 | 0 | 2,3 |
| CG9373 | FI21236p1 | 5 | 3 | 1 | 2,3 |
| CG2050 | DNA-binding protein modulo | 9 | 15 | 2 | 2,2 |
| CG11949 | Protein 4.1 homolog | 4 | 9 | 1 | 2 |
| CG13096 | Ribosomal L1 domain-containing protein CG13096 | 12 | 12 | 3 | 2 |
| CG8231 | GH13725p | 4 | 3 | 0 | 2 |
| CG2199 | FI17108p1 (Fragment) | 4 | 4 | 0 | 2 |
| CG1994 | RNA cytidine acetyltransferase | 4 | 2 | 0 | 2 |
| CG8977 | T-complex protein 1 subunit gamma | 4 | 1 | 0 | 2 |
| CG5374 | T-complex protein 1 subunit alpha | 4 | 2 | 0 | 2 |
| CG1345 | GH12731p | 4 | 1 | 0 | 2 |
| CG8351 | LD47396p | 4 | 1 | 0 | 2 |
| CG7033 | CG7033 | 4 | 0 | 0 | 2 |
| CG17521 | 60S ribosomal protein L10 | 4 | 6 | 0 | 2 |
| CG7283 | 60S ribosomal protein L10a-2 | 4 | 10 | 0 | 2 |
| CG9684 | LD24381p1 (Fragment) | 4 | 6 | 0 | 2 |
| CG3333 | MIP05689p (Fragment) | 11 | 4 | 3 | 1,9 |
| CG10922 | La protein homolog | 14 | 25 | 4 | 1,8 |
| CG10652 | FI02875p (Fragment) | 3 | 4 | 0 | 1,6 |
| CG1091 | Tailor, isoform C | 3 | 3 | 0 | 1,6 |
| CG1263 | 60S ribosomal protein L8 | 3 | 9 | 0 | 1,6 |
| CG9012 | Clathrin heavy chain | 3 | 2 | 0 | 1,6 |
| CG1883 | GM06992p | 3 | 1 | 0 | 1,6 |
| CG2033 | 40S ribosomal protein S15Aa | 3 | 1 | 0 | 1,6 |
| CG5525 | T-complex protein 1 subunit delta | 3 | 2 | 0 | 1,6 |
| CG10506 | Probable glutamine--tRNA ligase | 3 | 3 | 0 | 1,6 |
| CG30149 | Protein rigor mortis | 3 | 1 | 0 | 1,6 |
| CG4609 | Failed axon connections | 3 | 2 | 0 | 1,6 |

| | | | | | |
|---|---|---|---|---|---|
| CG5261 | Dihydrolipoamide acetyltransferase component of pyruvate dehydrogenase complex | 3 | 1 | 0 | 1,6 |
| CG9805 | Eukaryotic translation initiation factor 3 subunit A | 6 | 7 | 2 | 1,6 |
| CG4747 | Putative oxidoreductase GLYR1 homolog | 3 | 0 | 1 | 1,6 |
| CG4046 | 40S ribosomal protein S16 | 3 | 2 | 0 | 1,6 |
| CG42551 | La related protein, isoform F | 3 | 1 | 0 | 1,6 |
| CG4863 | LP14077p (Fragment) | 3 | 14 | 0 | 1,6 |
| CG7808 | 40S ribosomal protein S8 | 3 | 2 | 0 | 1,6 |
| CG7831 | Kinesin-like protein (Fragment) | 3 | 1 | 0 | 1,6 |
| CG7434 | 60S ribosomal protein L22 | 8 | 7 | 3 | 1,4 |
| CG5119 | Polyadenylate-binding protein | 12 | 14 | 5 | 1,3 |
| CG8280 | Elongation factor 1-alpha 1 | 11 | 8 | 5 | 1,1 |
| CG10686 | Trailer hitch, isoform G | 2 | 0 | 0 | 1 |
| CG10811 | Eukaryotic translation initiation factor 4G, isoform B | 2 | 1 | 0 | 1 |
| CG11276 | 40S ribosomal protein S4 | 2 | 1 | 0 | 1 |
| CG12775 | RE62581p | 2 | 5 | 0 | 1 |
| CG12785 | Nucleolar protein 6 | 2 | 2 | 0 | 1 |
| CG16901 | Squid, isoform E | 2 | 0 | 0 | 1 |
| CG2807 | RH74732p (Fragment) | 2 | 1 | 0 | 1 |
| CG3395 | 40S ribosomal protein S9 | 2 | 1 | 0 | 1 |
| CG3751 | 40S ribosomal protein S24 | 2 | 1 | 0 | 1 |
| CG4464 | 40S ribosomal protein S19a | 2 | 1 | 1 | 1 |
| CG4581 | Thiolase | 2 | 2 | 0 | 1 |
| CG3612 | ATP synthase subunit alpha, mitochondrial | 2 | 7 | 0 | 1 |
| CG4759 | 60S ribosomal protein L27 | 2 | 5 | 0 | 1 |
| CG4878 | Eukaryotic translation initiation factor 3 subunit B | 2 | 0 | 0 | 1 |
| CG14206 | RH14172p (Fragment) | 2 | 1 | 0 | 1 |
| CG8415 | 40S ribosomal protein S23 | 2 | 1 | 0 | 1 |
| CG11154 | ATP synthase subunit beta, mitochondrial | 2 | 3 | 0 | 1 |
| CG5920 | 40S ribosomal protein S2 | 2 | 1 | 0 | 1 |
| CG7961 | Coatomer subunit alpha | 2 | 1 | 0 | 1 |
| CG5520 | Glycoprotein 93 | 2 | 0 | 0 | 1 |
| CG12304 | Probable aminoacyl tRNA synthase complex-interacting multifunctional protein 2 | 2 | 1 | 0 | 1 |
| CG8439 | T-complex chaperonin 5, isoform B | 2 | 1 | 0 | 1 |
| CG5642 | Eukaryotic translation initiation factor 3 subunit L | 2 | 0 | 0 | 1 |
| CG4389 | Mitochondrial trifunctional protein alpha subunit, isoform B | 12 | 13 | 6 | 1 |
| | Histone H2B | 4 | 3 | 2 | 1 |
| | Histone H2A.v | 4 | 3 | 2 | 1 |
| CG6143 | Protein on ecdysone puffs, isoform D | 8 | 3 | 4 | 1 |
| CG6253 | GM06787p (Fragment) | 2 | 6 | 0 | 1 |
| | Histone H2A | 4 | 3 | 2 | 1 |
| CG9748 | Belle, isoform B | 6 | 3 | 3 | 1 |
| CG5170 | Dodeca-satellite-binding protein 1, isoform A | 4 | 5 | 2 | 1 |
| CG7726 | 60S ribosomal protein L11 | 2 | 4 | 1 | 1 |
| CG6510 | 60S ribosomal protein L18a | 2 | 5 | 0 | 1 |
| CG6779 | IP15838p (Fragment) | 2 | 0 | 0 | 1 |
| CG6846 | GEO07453p1 | 2 | 8 | 1 | 1 |
| CG7622 | 60S ribosomal protein L36 | 2 | 3 | 0 | 1 |
| CG8900 | FI09342p (Fragment) | 2 | 0 | 0 | 1 |
| | Histone H3 | 2 | 1 | 0 | 1 |
